# Public and private human T cell clones respond differentially to HCMV antigen when boosted by CD3 co-potentiation

**DOI:** 10.1101/2020.07.08.193805

**Authors:** Laura R.E. Becher, Wendy K. Nevala, Shari S. Sutor, Megan Abergel, Michele M. Hoffmann, Christopher A. Parks, Larry R. Pease, Adam G. Schrum, Svetomir N. Markovic, Diana Gil

**Affiliations:** Department of Immunology, Mayo Clinic Graduate School of Biomedical Sciences, Mayo Clinic, Rochester, MN, USA; Department of Oncology, Mayo Clinic, Rochester, MN, USA; Departments of Surgery and Molecular Microbiology & Immunology, School of Medicine, University of Missouri, Columbia, MO, USA; Department of Bioengineering, College of Engineering, University of Missouri, Columbia, MO, USA

## Abstract

Human cytomegalovirus (HCMV) induces long-lasting T cell immune responses that control but do not clear infection. Typical responses involve private T cell clones, expressing T cell antigen receptors (TCR) unique to a person, and also public T cell clones with identical TCRs active in different people. Here, we report the development of a pre-therapeutic immunostimulation modality against HCMV for human T cells, *CD3 co-potentiation*, and the clonal analysis of its effects in recall assays at single-cell resolution. CD3 co-potentiation of human T cells required identification of an intrinsically inert anti-CD3 Fab fragment that conditionally augmented signaling only when TCR was co-engaged with antigen. When applied in recall assays, CD3 co-potentiation enhanced the expansion of both public and private T cell clones responding to autologous HLA-A2(+) antigen-presenting cells and immunodominant NLV peptide from HCMV pp65 protein. Interestingly, public versus private TCR expression was associated with distinct clonal expansion signatures in response to recall stimulus. This implied that besides possible differences in their generation and selection in an immune response, public and private T cells may respond differently to pharmaco-immunomodulation. Furthermore, a third clonal expansion profile was observed upon CD3 co-potentiation of T cell clones from HLA-A2(-) donors and one HLA-A2(+) presumed-uninfected donor, where NLV was of low intrinsic potency. We conclude that human T cell copotentiation can increase the expansion of different classes of T cell clones responding to recall antigens of different strengths, and this may be exploitable for therapeutic development against chronic, persistent infections such as HCMV.

**Key Points:**

- Human CD3 co-potentiation can enhance the clonal expansion of several classes of recall T cells responding to antigens.
- Enhanced expansion follows a unique pattern based on the immunodominance or weakness of antigen, and public or private TCR status.

## Introduction

β-herpes human cytomegalovirus (HCMV) infects the population at high incidence^1^. Infection is often asymptomatic and controlled by long-lasting T cell responses driving the virus to latency, although sterilizing immunity is not induced^2–4^. Chronic inflammation or immune compromise caused by other infections, illnesses, aging, and drugs can allow HCMV reactivation and life-threatening disease^5–7^. Thus, there is great interest in developing new immune-boosting therapies to treat/prevent HCMV recurrence^8–10^.

In mice, we previously developed an immunostimulatory concept, *CD3 co-potentiation*, which, if applicable to humans, might add a new boosting strategy against a range of diseases from cancer to chronic viral infections, including HCMV. We described an anti-mouse-CD3 Mono-Fab fragment whose binding was functionally inert if T cells encountered no antigen, or if instead T cells were stimulated by strong antigens; but if T cells were stimulated by weak antigens, then coincident Fab:CD3 engagement improved various responses elicited from naïve CD8 T cells^11^. *In vivo,* the Fab reduced tumor burden of B16-F10 melanoma by a mechanism dependent on CD4 and CD8 T cells and TCR antigen specificity^11^. The anti-CD3 Fab induces a stimulation-poised CD3 conformation thought to amplify signaling upon weak antigen engagement by TCR^12,13^. Here, we report the development and characterization of an anti-human-CD3 Mono-Fab with co-potentiation function.

Whether CD3 co-potentiation can boost responses from previously activated (not naïve) T cells, and whether responses to strong antigens in addition to weak antigens might be enhanced are outstanding questions. HCMV(+) peripheral blood samples contain clonally expanded recall T cells that can be classified regarding their response to immunodominant versus weak HCMV antigens, and whether they bear public or private TCRs^14,15^. Here, we focus on immunodominant pp65^495-503^ peptide, NLVPMVATV (NLV) in HLA-A*02:01(+) individuals^16^, reported to induce both public and private CD8 T cell responses^17–24^. Both bulk recall assays and single-cell assessment of TCR clonality (TRBV CDR3 sequencing) from peripheral blood reveal that human CD3 co-potentiation can amplify expansion of public and private T cell clones. This effect occurs in response to (i) immunodominant NLV:HLA-A2, (ii) NLV as a weak antigen in HLA-A2(-) conditions and HLA-A2(+) presumed-uninfected conditions, and (iii) undefined HLA-dependent antigens in autologous antigen-presenting cells (APC). We propose that CD3 co-potentiation can amplify clonal expansion of recall T cells under various conditions of antigen experience and stimulatory strength, making the CD3 co-potentiation strategy potentially attractive for translation to HCMV and other disease treatments.

## Methods

### Cell lines

OT1ab.muCD8ab.JRT3 (OT-I.JRT3)^25^ and T2-Kb^12^ cells were reported previously. Both cell lines tested negative for mycoplasma and were grown in RPMI (Life Technologies) with 10% CosmicCalf serum (HyClone), 2 mM L-glutamine, and penicillin (100 U/ml)/streptomycin (100 μg/ml) (Life Technologies) at 37°C, 5% CO2.

### Peripheral blood mononuclear cells (PBMC)

With Mayo Institutional Review Board approval, whole human blood was collected from healthy volunteers. PBMCs were isolated by Ficoll gradient, harvested, washed, and used fresh or cryopreserved in FBS, 10% DMSO and stored in liquid nitrogen. Fresh or thawed PBMCs were washed and counted. Where indicated, T cells were isolated untouched using the magnetic human Pan-T-Cell Isolation Kit according to the manufacturer’s protocol (MACS, Miltenyi Biotec).

### Mono-Fab preparation

Mono-Fabs were purified as described previously^26^. Briefly, mAbs were digested by papain (Sigma-Aldrich) at 37°C for 24 hours. Digests were terminated with iodoacetamide (Sigma-Aldrich) and dialyzed in PBS at 4°C for 6 hours with periodic buffer exchange. Fc was removed via ProteinA Sepharose incubation (GE Healthcare) at 4°C overnight. Following further isolation by size exclusion chromatography over two tandem Superdex200 10/300GL columns (GE Life Sciences) on NGC-Quest10 FPLC system (BioRad), Mono-Fabs were sterile-filtered and maintained in cold PBS + 2M L-proline to preserve monovalency^27^. Protein concentration was quantified on DeNovixDS-11 Spectrophotometer.

### Peptides and antibodies

FARL (SSIEFARL; no OT-I-TCR specificity) and OVA (SIINFEKL) peptides were purchased from Elim Biopharmaceuticals. NLV peptide (NLVPMVATV) was synthesized in the Proteomics Core at Mayo Clinic. OKT3-producing hybridoma was kindly provided by Ed Palmer (University of Basel, Switzerland) and purified in-house from hybridoma supernatant. UCHT1 mAb was purchased from BioXcell. Anti-CD8 DK25 mAb was purchased from Agilent. Serum Ms-IgG and Ms-IgG-Fab were purchased from Jackson ImmunoResearch. Fluorescent conjugated antibodies used for flow cytometry included: anti-mouse IgG (BioLegend, Poly4060), anti-human CD4 (BD Biosciences, RPA-T4, OKT4), anti-human CD8 (BD Biosciences, HIT8a, RPA-T8, SK1), anti-human CD69 (BD Biosciences, FN50), anti-human CD45 (BioLegend, UCHL1), anti-human CD56 (BD Biosciences, B159), anti-human CD19 (BioLegend, HIB19), anti-Vβ5 (BioLegend, MR9-4), and anti-Nur77 (Invitrogen, 12.14). H2-Kb/OVA-tetramer was made in house as previously described^28^. HLA-A*02:01/NLV (A2/NLV)-tetramer was purchased from MBL International. GhostDye discriminated live/dead cells (Tonbo).

### Flow cytometry

Cells were stained with indicated fluorophore-conjugated antibodies and samples were collected on either Guava easyCyte HT Flow Cytometer (Luminex) or BD-Accuri C6 Flow Cytometer (BD Biosciences). For intracellular staining of Nur77, samples were fixed and permeabilized with Cytofix/Cytoperm Kit per the manufacturer’s protocol (BD Biosciences). Data analysis was performed using FlowJo (Tree Star) or guavaSoft software.

### T cell activation

To measure OT-I.JRT3 T cell responses, 50,000 cells/well were stimulated with 0.2nM indicated peptides presented by T2-Kb APCs and 10μg/mL Ms-IgG-Fab control or specific Mono-Fabs for 24 hours, followed by flow cytometry analysis of CD69 upregulation and TCR downregulation. For primary human T cell responses, PBMCs were rested overnight following fresh isolation or thaw from cryopreservation. PBMCs were seeded at 0.2×10^6^ cells per well in a round bottom 96-well culture plate and stimulated with indicated control or specific immunoglobulins (10μg/mL) for 4-6 hours, followed by flow cytometry analysis of CD69 and Nur77 upregulation.

### Western blot

1×10^6^ human PBMCs were stimulated with 10μg/mL immunoglobulins as indicated, in the presence or absence of pervanadate for 5 minutes, 37°C. Cells were washed twice in cold PBS and lysed for 10 minutes in 1% TritonX-100, 20mM Tris/HCl pH 7.4, 150mM NaCl, plus Halt-protease/phosphatase inhibitors (ThermoFisher). Cells were pelleted by centrifugation and equivalent cell lysates were subjected to SDS-PAGE (reducing, 10% gel), PVDF membrane transfer, and Western blot analysis with anti-phosphotyrosine (EMD Millipore, 4G10) and secondary anti-mouse IgG HRP (Cell Signaling).

### CD3 pull-down (CD3-PD)

The CD3-PD assay was used to quantify CD3 conformational change (CD3Δc) by detection of CD3ε proline-rich sequence exposure^11–13,29^. Briefly, 30×10^6^ PBMCs from healthy human donors were lysed in isotonic buffer containing 1% Brij58 (Sigma) on ice for 30 minutes, followed by centrifugation to obtain post-nuclear fractions. Samples were pre-cleared with GST beads (4°C, 1 hour) in the presence of indicated immunoglobulins (10μg/mL), followed by specific CD3-PD with GST-SH3.1-NCK beads (4°C, 12 hours). CD3-PD samples were subjected to SDS-PAGE (reducing, 13% gel), nitrocellulose transfer, and Western blot analysis with rabbit serum 448 antibody, specific for CD3ζ (kind gift from Balbino Alarcón, Universidad Autónoma de Madrid, Madrid, Spain). The mAb APA 1/1 (GE Biosciences) set the assay background level^29^. Protein acetone-precipitates from a fraction of post-nuclear lysates controlled for total CD3 content per sample, and quantification was performed as described previously^13^.

### Recall T cell expansion cultures

CD8 T cells were expanded as described previously^30^ with minor modifications. Total PBMCs were seeded on Day 0 at 0.1-0.2×10^6^ cells per well in round bottom 96-well culture plates with no peptide or 1uM NLV added in RPMI, 10% FBS. On Day 2, 10ug/mL Mono-Fab and 20 U/mL IL-2 (Proleukin, Mayo Pharmacy) were added to culture. On Days 4 and 7, half the media was replaced with fresh media containing 20 U/mL IL-2. Flow cytometry was run on Day 9. For antigen blocking experiments, 5ug/mL DK25 antibody or Ms-IgG control was added on Days 0, 2, and 4, and flow cytometry was run on Day 7.

### TRBV CDR3 sequencing and analysis

Genomic DNA was isolated from frozen cell pellets (QIAamp DNA mini kit). TRBV-CDR3 sequencing and preliminary analysis was completed using the ImmunoSEQ platform^31^ (Adaptive Biotechnologies, hsTCRβ kit). Per the manufacturer’s protocol, 1.6 μg of genomic DNA per sample was subjected to PCR to amplify all TRBV-CDR3 sequences in a bias-controlled manner using multiplexed V- and J-gene primers. Amplified TRBV-CDR3 underwent a second PCR to generate barcoded libraries. Sequencer-ready barcoded libraries were pooled and sequenced on an Illumina MiSeq with primers provided in the hsTCRβ kit. Raw sequencing data was sent to Adaptive Biotechnologies for processing to report the normalized, annotated TCRβ repertoire of each sample. Data analysis was performed using the provided immunoSEQ Analyzer program. The free VDJdb database^32^ was accessed to identify TRBV CDR3 clones in the dataset matching those from previously published public TCRs associated with HLA-A2 and NLV peptide.

### Statistical analysis

Statistics performed using GraphPad Prism included two-tailed, one-tailed, unpaired, and paired Student’s *t*-tests and Fischer’s Exact Test. Results showing central values represent mean ± SD or SEM.

### Data sharing statement

For original data, please contact.

## Results

### Mono-OKT3-Fab binds to human CD3 without blocking TCR:antigen interactions

Two anti-human-CD3ε Fab fragments were prepared by papain digestion of their corresponding mAbs, OKT3 and UCHT1. Digestion was followed by removal of Fc-domain-containing fragments and size exclusion chromatography to purify Fabs as monovalent species of ~50 kDa. Critically, to maintain monovalency and prevent auto-dimerization, Fabs were stored in the presence of the osmolyte L-proline and are referred to as Mono-Fabs to emphasize careful control of monovalency during their production and storage^26^. For co-potentiation, Mono-Fabs must bind CD3 without sterically hindering TCR:antigen binding or signaling. OKT3 and UCHT1 Mono-Fabs bound surface CD3 of OT-I.JRT3 cells, expressing mouse CD8αβ and OT-I-TCRαβ in complex with human CD3 ^25^ (**Figure 1A**, left). Both Mono-Fabs also bound primary human CD4 and CD8 T cells (**Figure 1A**, middle and right). When bound, Mono-OKT3-Fab did not block TCR:antigen interaction, demonstrated by unaffected Kb/OVA-tetramer staining of OT-I.JRT3 cells (**Figure 1B**, top left), and uninterrupted A2/NLV-tetramer staining of HLA-A*02:01(+) CD8 T cells (**Figure 1B**, top right). Furthermore, binding of Mono-OKT3-Fab to OT-I.JRT3 cells did not alter surface TCR downregulation or CD69 upregulation in response to SIINFEKL antigenic peptide (**Figure 1C**, pOVA). In contrast, Mono-UCHT1-Fab inhibited TCR:antigen interaction in both OT-I.JRT3 and human A2/NLV-tetramer(+) CD8 T cells (**Figure 1B**, bottom) and inhibited surface TCR downregulation and CD69 upregulation of OT-I.JRT3 cells in response to SIINFEKL (**Figure 1C**, pOVA).

**Figure 1.**
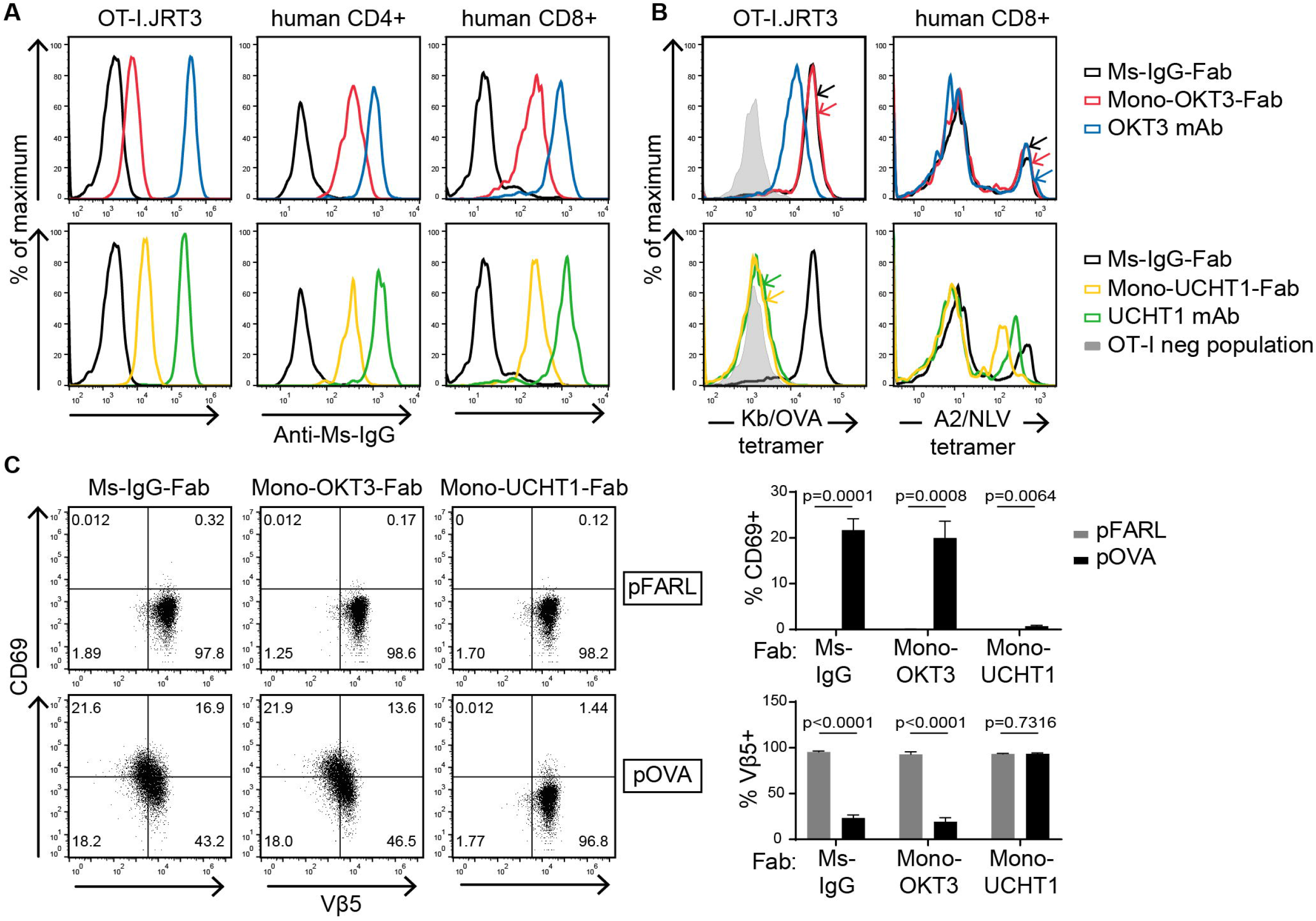
Mono-OKT3-Fab binds to human T cells without blocking TCR:antigen interactions. (**A**) Mono-OKT3-Fab and Mono-UCHT1-Fab bind T cells, detected by positive secondary anti-Ms-IgG staining by flow cytometry of OT-I.JRT3 cells and primary human CD4 and CD8 T cells from PBMCs. (**B**) Mono-OKT3-Fab does not block TCR:antigen binding in contrast to Mono-UCHT1-Fab. OT-I.JRT3 or CD8 T cells that were previously expanded with NLV peptide were pre-incubated with indicated Igs and stained for binding of Kb/OVA tetramer (B, left) or A2/NLV tetramer (B, right), respectively. (**C**) Mono-OKT3-Fab does not impair the T cell response to cognate antigen, unlike Mono-UCHT1-Fab. OT-I.JRT3 cells were cultured with null peptide (pFARL) or antigenic peptide (pOVA) presented on T2-Kb APCs in the presence of indicated Igs and analyzed for CD69 upregulation and TCR downregulation. Frequencies of CD69(+) and Vβ5(+) cells are shown (mean ± SD from triplicate samples, two-tailed unpaired Student’s t-test). Panels (A-C) are representative of three or more independent experiments. Ms-IgG-Fab, negative control.

### Mono-OKT3-Fab induces CD3Δc without initiating early signaling

In the absence of antigen recognition, neither Mono-Fab induced signaling-dependent surface TCR downregulation or CD69 upregulation in OT-I.JRT3 cells, as expected for non-crosslinking species (**Figure 1C,** pFARL). Likewise, neither Mono-Fab induced CD69 or Nur77 upregulation in primary human T cells, unlike their parent bivalent mAbs (**Figure 2A**). Furthermore, Mono-Fabs did not induce accumulation of tyrosine-phosphorylated proteins following engagement of human PBMCs compared to positive control, pervanadate^33^ (**Figure 2B**). Despite their inability to trigger CD3 signaling, Mono-Fab-binding induced a signal-amplifying conformational change in CD3 (CD3Δc), indicated by a CD3-PD assay (**Figure 2C**), where GST-SH3.1-Nck beads capture TCR/CD3 complexes displaying a CD3ε proline-rich sequence exposed upon optimal TCR engagement^11–13,29^. Based on these results, Mono-OKT3-Fab was selected to study co-potentiation of human T cells due to its ability to bind human CD3 and induce CD3Δc without blocking TCR:antigen interaction and without intrinsically initiating early signaling.

**Figure 2.**
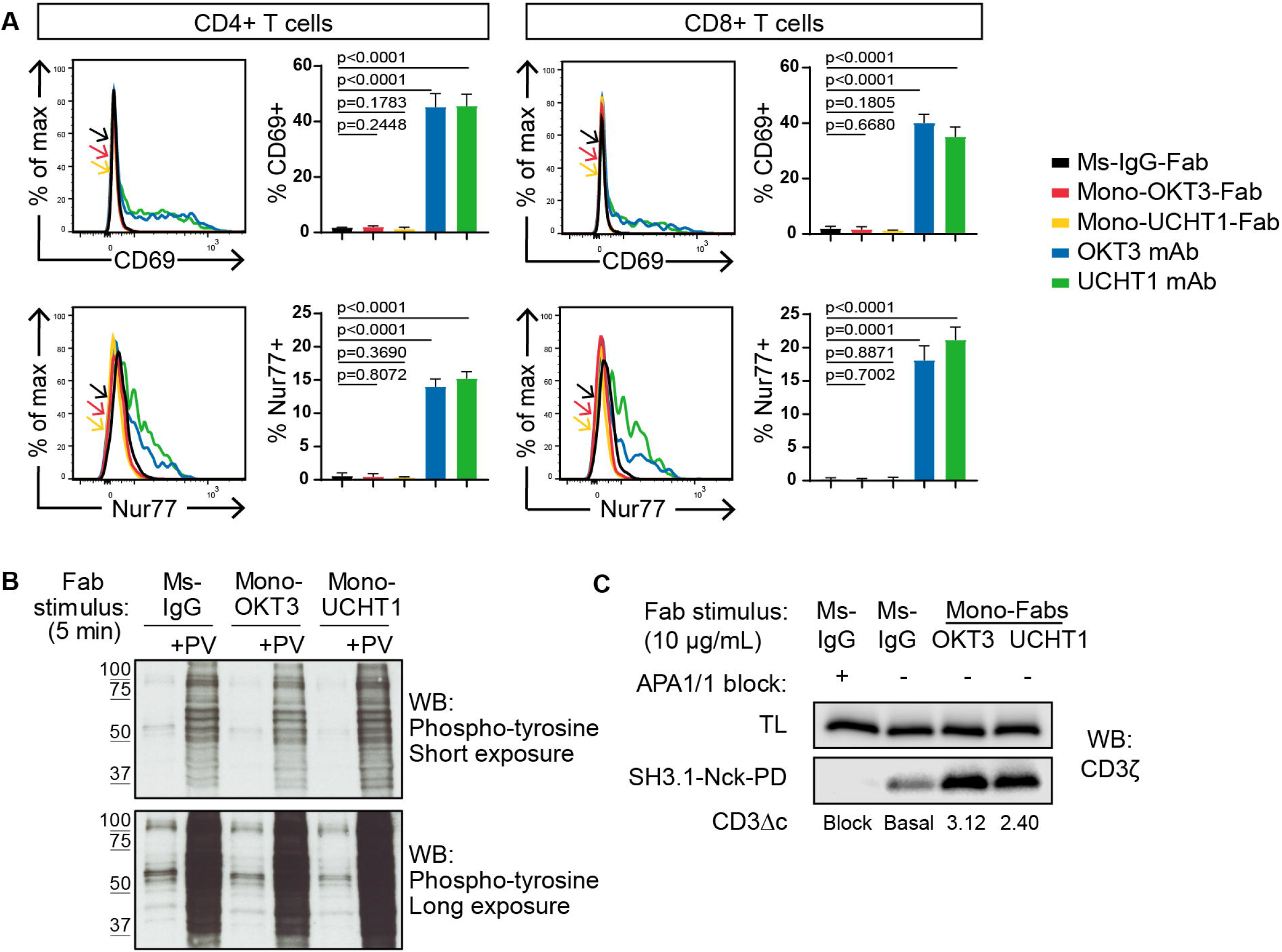
Mono-Fabs induce CD3Δc in the human TCR/CD3 complex without initiating early T cell signaling. (**A-B**) Binding of Mono-Fabs does not stimulate T cells in the absence of antigen. (A) PBMCs were incubated with indicated Igs, after which CD4 and CD8 T cells were analyzed for the induction of surface CD69 and intracellular Nur77 by flow cytometry. Frequencies of CD69(+) and Nur77(+) T cells are shown (mean ± SD from triplicate samples, two-tailed unpaired Student’s t-test). (B) PBMCs were incubated with indicated Igs in the presence or absence of pervanadate (PV). Phosphotyrosine was detected by Western blot (WB) of equivalent cell lysates. (**C**) Mono-Fabs induce CD3Δc. PBMC lysates were incubated with APA1/1 (to block CD3 pull-down), Ms-IgG-Fab (to reveal basal level of CD3Δc), or Mono-Fabs (test conditions) and assessed for induction of CD3Δc by the CD3 pull-down assay. Post- CD3Δc open conformation was detected with anti-CD3ζ by WB. Inducible CD3Δc is measured by fold-increase over basal level. TL, total lysate before pull-down. Panels (A-C) are representative of three or more independent experiments.

### Mono-OKT3-Fab enhances recall T cell expansion to autologous APCs and NLV:HLA-A2 by a mechanism dependent on TCR:HLA and CD8 co-receptor engagement

Seven healthy blood donors were classified by HLA-A*02:01 expression and the presence of CD8 T cells positive for binding A2/NLV-tetramer, an immunodominant antigen and marker of HCMV positivity. Four donors were HLA-A*02:01(+) and A2/NLV-tetramer(+) (72F, 53M, 28M, 47M), one donor was HLA-A*02:01(+) but A2/NLV-tetramer(-) (74M), and two donors were HLA-A*02:01(-) and A2/NLV-tetramer(-) (78F, 59F; **Supplemental Table 1**). After culturing PBMCs for nine days in the presence of exogenous NLV + irrelevant Ig + late-IL-2 (see Methods), donors positive for HLA-A*02:01 and A2/NLV-tetramer presented higher CD8 T cell counts than cultures without exogenous peptide (**Figure 3A**, gray). These donors were categorized as <u>exogenous NLV-responsive in bulk culture (exog-NLV-bulk-responsive</u>, **Supplemental Table 1**). Cell number increased even more in the presence of Mono-OKT3-Fab (**Figure 3A**, red). In contrast, A2/NLV-tetramer(-) cells in the same cultures increased in the presence of Mono-OKT3-Fab, but not exogenous peptide (**Figure 3B**), and PBMCs from A2/NLV-tetramer(-) donors responded likewise (**Figure 3C**). These data show that Mono-OKT3-Fab increased the numbers of both A2/NLV-tetramer(+) and A2/NLV-tetramer(-) CD8 T cells.

**Figure 3.**
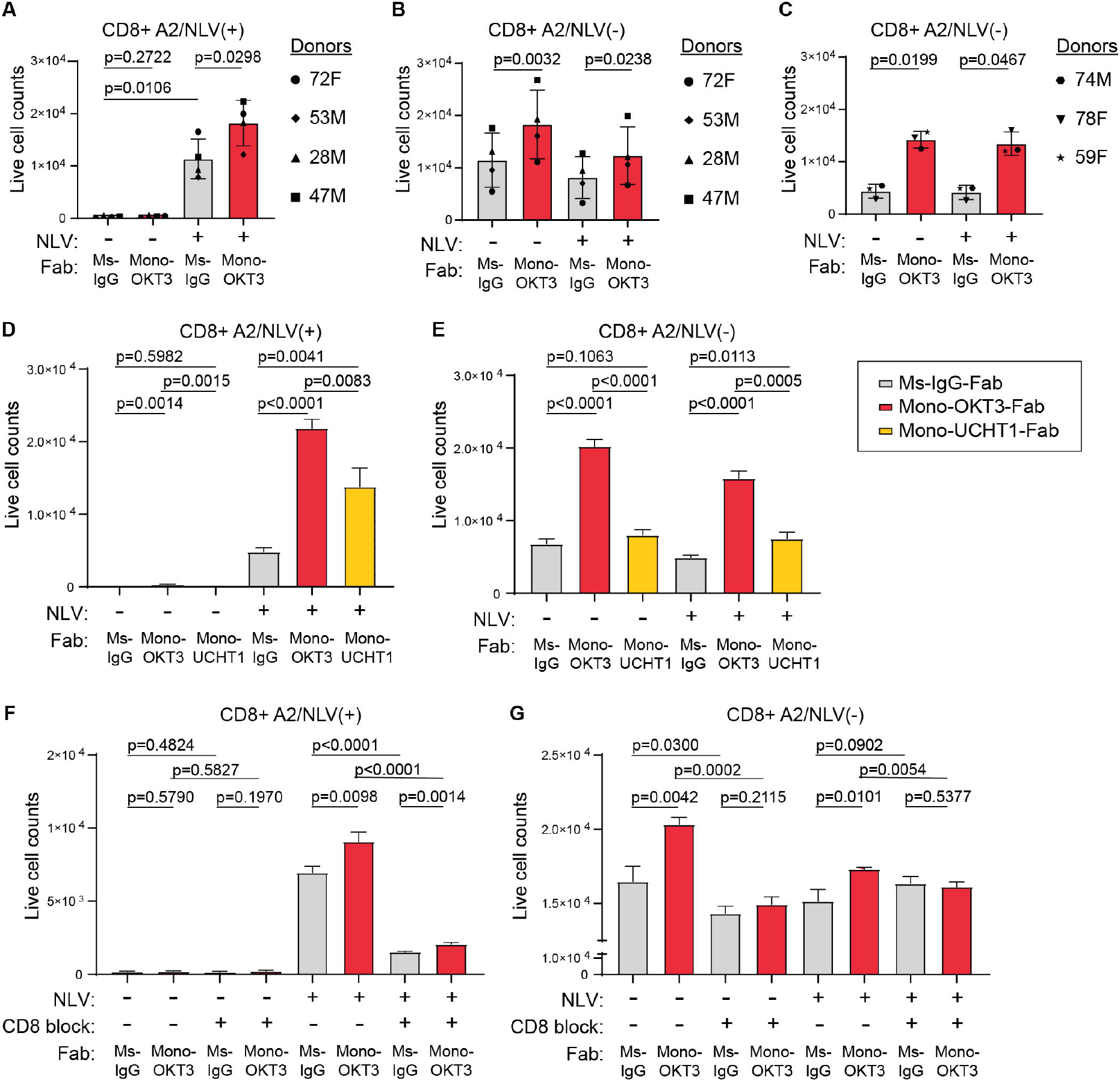
Mono-OKT3-Fab enhances recall T cell expansion to NLV:HLA-A2 or autologous APCs- only by a mechanism dependent on TCR:HLA and CD8 co-receptor engagement. (A-C) PBMCs were cultured +/− exogenous NLV peptide in the presence of Mono-OKT3-Fab or control Ms-IgG-Fab for 9 days. (**A**) On Day 9, cells were analyzed by flow cytometry for the number of expanded A2/NLV- tetramer(+) CD8 T cells from exog-NLV-bulk-responsive donors. In combination with exogenous NLV, Mono-OKT3-Fab increased the number of A2/NLV-tetramer(+) CD8 T cells. (**B-C**) Mono-OKT3-Fab also increased the number of A2/NLV-tetramer(-) CD8 T cells in exog-NLV-bulk-responsive donors (B) and in exog-NLV-bulk-non-responsive donors (C). For (A-C), each symbol represents the average of at least three independent experiments per donor (mean ± SD, two-tailed paired Student’s t-test). (**D-E**) PBMCs were cultured +/− exogenous NLV peptide in the presence of Ms-IgG-Fab (control), Mono- OKT3-Fab, or Mono-UCHT1-Fab for 9 days. Mono-UCHT1-Fab dampened the co-potentiation effect as compared to Mono-OKT3-Fab in both A2/NLV-tetramer(+) (D) and A2/NLV-tetramer(-) (E) CD8 T cells. (**F-G**) PBMCs were cultured +/− exogenous NLV peptide in the presence of Ms-IgG-Fab or Mono- OKT3-Fab and +/− the CD8 blocking antibody, DK25, for 7 days. Blocking CD8 reduced the copotentiation effect of Mono-OKT3-Fab for both A2/NLV-tetramer(+) (F) and A2/NLV-tetramer(-) (G) CD8 T cells. For (D-G), one representative experiment of Donor 47M is shown of three replicates (mean ± SD from triplicate samples, two-tailed unpaired Student’s t-test).

To distinguish Mono-OKT3-Fab intrinsic T cell stimulation from TCR:HLA-dependent copotentiation, recall expansion assays were performed in the presence or absence of blocking reagents to TCR:HLA or CD8 co-receptor. First, CD8 T cells from one exog-NLV-bulk-responsive donor were cultured in the presence of Mono-OKT3-Fab or Mono-UCHT1-Fab, the latter binding to CD3 and inducing CD3Δc but impairing antigen binding to T cells (**Figures 1–2**). Mono-UCHT1-Fab significantly reduced co-potentiation when compared with Mono-OKT3-Fab of CD8 T cells either positive (**Figure 3D**) or negative (**Figure 3E**) for A2/NLV-tetramer. These results indicate that with impaired TCR:antigen interactions, induction of CD3Δc by Mono-Fabs is insufficient to mediate co-potentiation. Second, parallel experiments showed that the anti-CD8 blocking antibody, DK25^34^, inhibited both exogenous-NLV-specific and Mono-OKT3-Fab-specific responses in CD8 T cells either positive (**Figure 3F**) or negative (**Figure 3G**) for A2/NLV-tetramer. Together, these data indicate that Mono-OKT3-Fab copotentiation is dependent on CD8:TCR:HLA engagement, for both A2/NLV-tetramer(+) CD8 T cells driven by exogenous NLV and for A2/NLV-tetramer(-) CD8 T cells driven only by autologous APCs.

### Mono-OKT3-Fab co-potentiation primarily enhances expansion of top-ranked T cell clones

To analyze the T cell clonal dynamics of the co-potentiation response, recall assays were followed by DNA extraction and TRBV-CDR3 analysis via ImmunoSeq with single-cell resolution (Adaptive Technologies, see Methods). The number of clones sampled per donor per culture condition reached as high as ~50,000. Clonal diversity was estimated by scaled Shannon entropy, a value whose range is 0-1, where 0 represents minimum diversity exhibited by a monoclonal T cell population and 1 represents maximal repertoire diversity when all TRBV-CDR3 sequences are expressed equally^35^. We observed that exogenous NLV decreased entropy compared with negative-control cultures for 4/4 exog-NLV-bulk-responsive donors, and likewise decreased the total clone number sampled from cultures (**Supplemental Figure 1A-H**), an expected outcome when T cell clones specific for a single peptide proliferate and increase their relative representation. In contrast, Mono-OKT3-Fab did not reliably produce such an effect (observed in 2/4 exog-NLV-bulk-responsive donors) nor did Mono-OKT3-Fab tend to further decrease entropy when administered in combination with exogenous NLV compared with NLV alone (observed in 1/4 exog-NLV-bulk-responsive donors; **Supplemental Figure 1A-H**). Thus, Mono-OKT3-Fab was not producing a clonal effect identical to that of exogenous peptide.

To determine if Mono-OKT3-Fab indiscriminately caused many clones to expand, we used single-cell clonal sequencing data to estimate total copy number of each clone in recall cultures and compared between conditions by rank analysis. Interestingly, we found that despite thousands of clones measured, substantial differences in cell numbers ranked according to abundance were heavily concentrated in the top-ten clones (**Figure 4A-D; Supplemental Figure 1I-L**). Thus, co-potentiation must have a mechanism of clonal specificity, which is more deeply analyzed here, discussing the HLA-A2+ exog-NLV-bulk-responsive Donor 72F as an example. Among the top-ten clones from *negativecontrol* cultures, none were NLV-responsive (**Supplemental Table 2, Donor 72F**; no exogenous peptide+Ms-IgG-Fab, rows 4-13), and most scored as “zeros” in the other culture conditions. This likely indicates the individual clones were too infrequent for consistent sampling. In *exogenous NLV* cultures, the top clone reached ~130,000 cells (**Supplemental Table 2, Donor 72F**; exogenous NLV peptide+Ms-IgG-Fab, rows 16-25), having represented ~1500 cells in negative-control culture. Several other top clones were absent in one or more other culture conditions and thus, as above, some analysis had sampling limitations. However, ranks 3 and 5 were consistently sampled (**Supplemental Table 2, Donor 72F**; no exogenous peptide+Mono-OKT3-Fab, rows 18 and 20) and showed matching clones that were also amplified in the *Mono-OKT3-Fab-only* condition, with synergistic highest abundance in NLV+Mono-OKT3-Fab cultures. Rank 8 was already high in negative-control culture and was neither exogenous-NLV-responsive nor amplified by Mono-OKT3-Fab (**Supplemental Table 2, Donor 72F**; no exogenous peptide+Mono-OKT3-Fab, row 23). In Mono-OKT3-Fab-only cultures (**Supplemental Table 2, Donor 72F**; no exogenous peptide+Mono-OKT3-Fab, rows 28-37), 3/10 top clones were independently NLV-responsive in exogenous-NLV cultures, while 4/10 top clones responded to Mono-OKT3-Fab but not exogenous NLV. Finally, in *NLV+Mono-OKT3-Fab* cultures (**Figure 4E** and **Supplemental Table 2, Donor 72F**; exogenous NLV peptide+Mono-OKT3-Fab, rows 40-49), 4/10 top clones were NLV-responsive and synergistically amplified, while 5/10 top clones were not NLV-responsive but were amplified to a similar extent as Mono-OKT3-Fab-only cultures. Therefore, copotentiation amplified clonal abundance of certain top clones only, with some but not others also being responsive to exogenous NLV.

**Figure 4.**
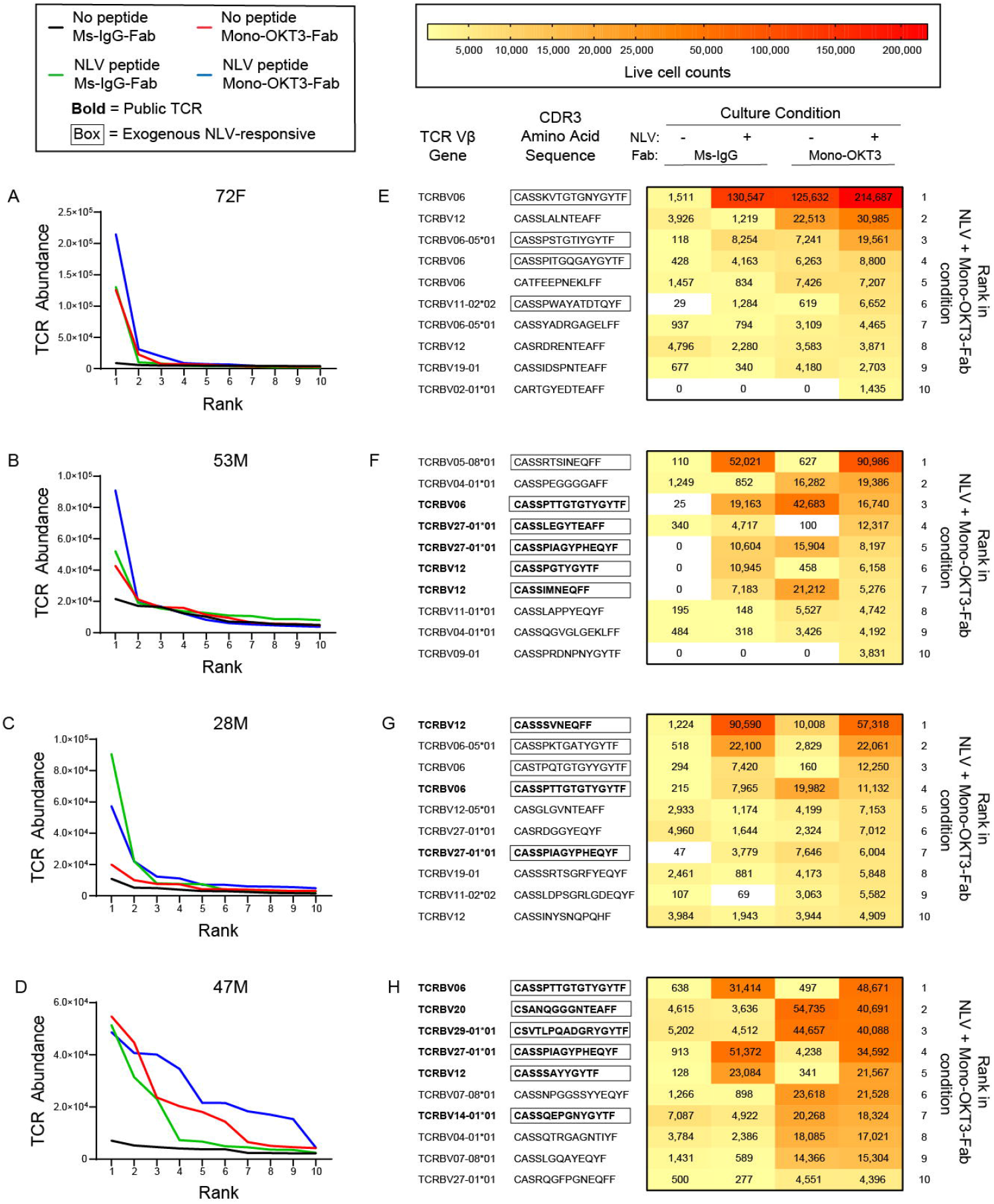
Mono-OKT3-Fab co-potentiation primarily enhances expansion of top-ranked T cell clones. (**A-D**) TCR clones for each exog-NLV-bulk-responsive donor were ranked from most abundant to least abundant for each condition. Differences in rank-versus-rank performance concentrated in the top-10 clones. (**E-H**) Among top ranked clones, there was variability in the extent to which clones were amplified by exogenous NLV, Mono-OKT3-Fab, or both in combination. The top-10 ranked clones from the NLV+Mono-OKT3-Fab condition are shown with their corresponding live cell number abundance in the other three conditions. Boxed amino acid sequences indicate NLV-specific clones (clones with greater abundance in NLV+Ms-IgG-Fab versus No peptide+Ms-IgG-Fab condition, or for public TCRs, observation of that pattern in at least one other donor or previously reported in the literature). Sequences in bold represent public TCR-bearing clones appearing in multiple donors in the present study, or previously reported in the literature. Heatmaps visualize the increase in clonal cell number generated by exogenous NLV, Mono-OKT3-Fab, or both in combination.

### Different classes of T cell clones respond to CD3 co-potentiation with distinct clonal expansion signatures

The other three exog-NLV-bulk-responsive donors showed similar examples of clonal responses (**Figure 4F-H**), while unlike Donor 72F, the others had exogenous-NLV-responsive public TCRs (**Supplemental Table 3A**) among top clones, which in 12/14 occurrences responded to Mono-OKT3-Fab (**Supplemental Table 3B**). However, there was a curious pattern in their response: public clones tended to respond best to either Mono-OKT3-Fab-only or exogenous NLV, but less optimally to the combination. Assigning first-place performance “Gold” to conditions with highest clonal abundance, and second/third-place “Silver-Bronze”, we found that considering all top-ten clones, private much more than public TCRs showed synergy in combination treatment (**Table 1**).

**Table 1.**
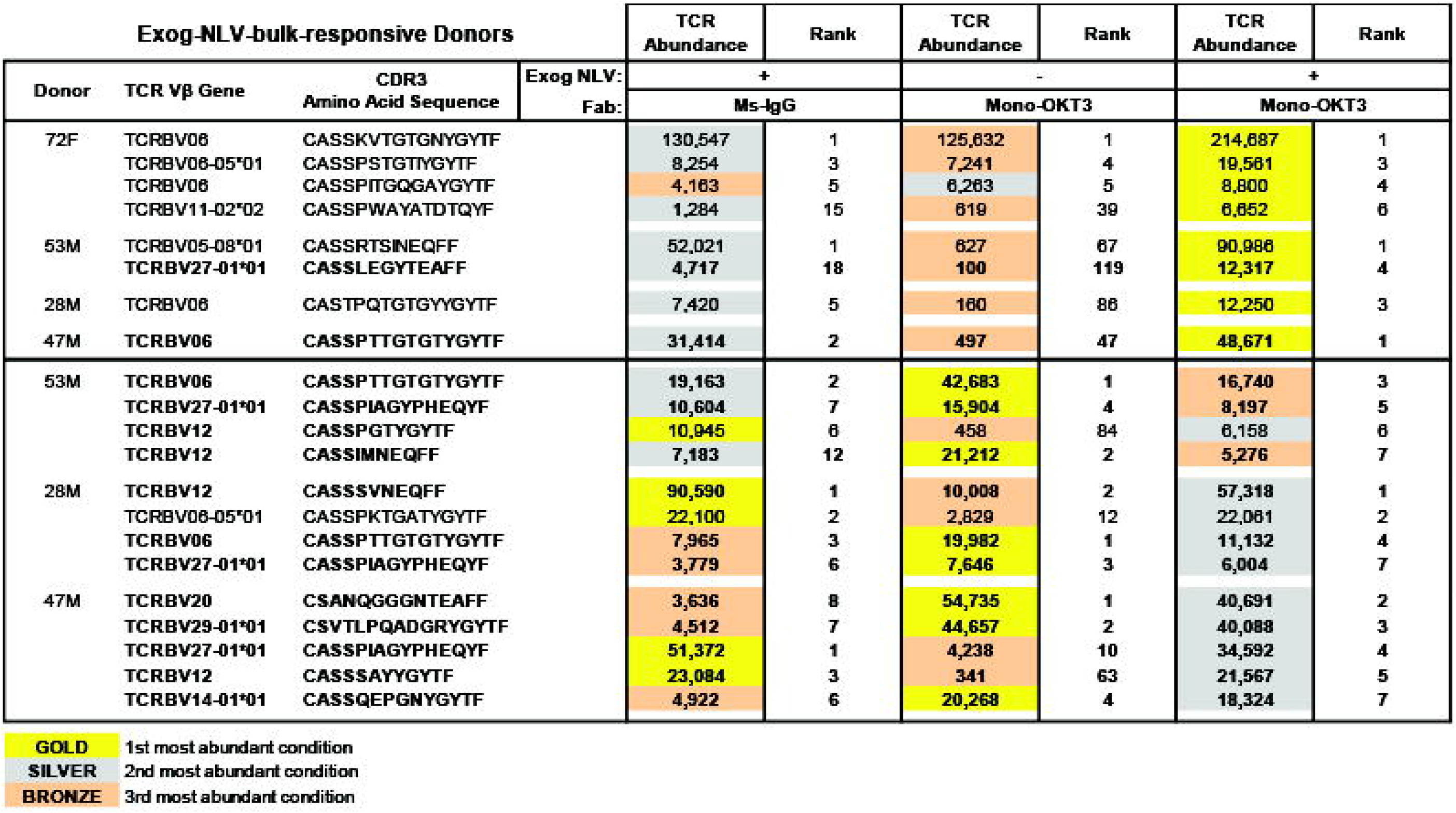
In exog-NLV-bulk-responsive donors, private TCRs tend to show synergy in combination treatment while public TCRs respond best to either Mono-OKT3-Fab or exogenous NLV separately. TCR clones ranked in the top-10 of NLV+Mono-OKT3-Fab condition with evidence of NLV-specificity were analyzed according to their abundance in various recall culture conditions. For each TRBV CDR3 amino acid sequence, “Gold” was awarded to conditions with the highest clonal abundance, “Silver” to second place, and “Bronze” to third place. Bolded sequences indicate public TCRs while the others are private TCRs. Evidence for NLV-specificity was accepted as displaying higher clone numbers in NLV+Ms-IgG-Fab versus No peptide+Ms-IgG-Fab conditions, or, for public TCRs, that pattern in at least one other donor or previously reported in the literature. The tendency for public TCRs to score Gold in NLV-only or Mono-OKT3-Fab-only treatments and private TCRs to score Gold in combination treatment was statistically significant (p = 0.003, two-tailed Fischer’s Exact Test; p = 0.007, chi-square test with Yates correction).

| Exog-NLV-bulk-responsive Donors |  |  |  | TCR Abundance | Rank | TCR Abundance | Rank | TCR Abundance | Rank |
| --- | --- | --- | --- | --- | --- | --- | --- | --- | --- |
| Donor | TCR Vβ Gene | CDR3 Amino Acid Sequence | Exog NLV: Fab: | + |  | - |  | + |  |
|  |  |  |  | Ms-IgG |  | Mono-OKT3 |  | Mono-OKT3 |  |
| 72F | TCRBV08 | CASSKVTGTGNYGYTF |  | 130,547 | 1 | 125,632 | 1 | 214,687 | 1 |
|  | TCRBV08-05*01 | CASSPSTGTIYGYTF |  | 8,254 | 3 | 7,241 | 4 | 19,561 | 3 |
|  | TCRBV08 | CASSPITGQGAYGYTF |  | 4,163 | 5 | 6,263 | 5 | 8,800 | 4 |
|  | TCRBV11-02*02 | CASSPWAYATDTQYF |  | 1,284 | 15 | 619 | 39 | 6,652 | 6 |
| 53M | TCRBV05-08*01 | CASSRTSINEQFF |  | 52,021 | 1 | 627 | 67 | 90,986 | 1 |
|  | TCRBV27-01*01 | CASSLEGYTEAFF |  | 4,717 | 18 | 100 | 119 | 12,317 | 4 |
| 28M | TCRBV08 | CASTPQTGTGYGYTF |  | 7,420 | 5 | 160 | 86 | 12,250 | 3 |
| 47M | TCRBV06 | CASSPTTGTGTGYGYTF |  | 31,414 | 2 | 497 | 47 | 48,671 | 1 |
| 53M | TCRBV06 | CASSPTTGTGTGYGYTF |  | 19,163 | 2 | 42,683 | 1 | 16,740 | 3 |
|  | TCRBV27-01*01 | CASSPIAGYPHEQYF |  | 10,604 | 7 | 15,904 | 4 | 8,197 | 5 |
|  | TCRBV12 | CASSPGTYGYTF |  | 10,945 | 6 | 458 | 84 | 6,158 | 6 |
|  | TCRBV12 | CASSIMNEQFF |  | 7,183 | 12 | 21,212 | 2 | 5,276 | 7 |
| 28M | TCRBV12 | CASSSVNEQFF |  | 90,590 | 1 | 10,008 | 2 | 57,318 | 1 |
|  | TCRBV08-05*01 | CASSPKTGATYGYTF |  | 22,100 | 2 | 2,829 | 12 | 22,061 | 2 |
|  | TCRBV06 | CASSPTTGTGTGYGYTF |  | 7,965 | 3 | 19,982 | 1 | 11,132 | 4 |
|  | TCRBV27-01*01 | CASSPIAGYPHEQYF |  | 3,779 | 6 | 7,646 | 3 | 6,004 | 7 |
| 47M | TCRBV20 | CSANQGGGNTAEFF |  | 3,636 | 8 | 54,735 | 1 | 40,691 | 2 |
|  | TCRBV29-01*01 | CSVTLPPQADGRYGYTF |  | 4,512 | 7 | 44,657 | 2 | 40,088 | 3 |
|  | TCRBV27-01*01 | CASSPIAGYPHEQYF |  | 51,372 | 1 | 4,238 | 10 | 34,592 | 4 |
|  | TCRBV12 | CASSSAYYGYTF |  | 23,084 | 3 | 341 | 63 | 21,567 | 5 |
|  | TCRBV14-01*01 | CASSQEPGNYGYTF |  | 4,922 | 6 | 20,268 | 4 | 18,324 | 7 |

|  |  |
| --- | --- |
| **GOLD** | 1st most abundant condition |
| **SILVER** | 2nd most abundant condition |
| **BRONZE** | 3rd most abundant condition |

We next examined the T cell clonal dynamics of co-potentiation in the three A2/NLV-tetramer(-) donors, all of which amplified T cell expansion by Mono-OKT3-Fab but were <u>exog-NLV -bulk-non-responsive</u> (**Figure 3C**). Exogenous NLV and Mono-OKT3-Fab status did not correlate with predictable changes in entropy and total clones sampled, although rank differences remained largely concentrated in top-ten clones (**Supplemental Figure 2**). Interestingly, the top clone for each donor appeared exogenous-NLV-responsive, as did several other top clones, with further enhancement combined with Mono-OKT3-Fab (**Supplemental Table 4**). To assess NLV+Mono-OKT3-Fab combinatorial synergy, we applied Gold/Silver-Bronze analysis to these 13 apparently NLV-responsive clones, and found that in all cases

NLV+Mono-OKT3-Fab condition produced the highest clonal abundance (**Figure 5A**). Despite this similarity with the “private TCR” signature noted previously for exog-NLV-bulk-responsive donors, there was also a distinct difference. The exogenous-NLV-only condition always but barely increased the cell number of these clones above the negative-control culture condition (~1-4-fold); in contrast, combination-treatment “Gold” response clones from exog-NLV-bulk-responsive donors were much more peptide-responsive, (~10-500-fold, **Figure 5B; Supplemental Table 5**). We also found that comparing clonal cell numbers from cultures with Mono-OKT3-Fab +/− exogenous NLV, combination-treatment “Gold” response clones from exog-NLV-bulk-responsive donors appeared in two clusters: one for which exogenous NLV peptide increased clonal abundance by ~75-150-fold, and another which only increased ~1-11-fold; in contrast, NLV-responsive clones from exog-NLV-bulk-non-responsive donors all appeared in the low-peptide-response cluster (**Figure 5C**). This pattern flipped when assessing the contribution of Mono-OKT3-Fab to combinatorial synergy: here, exog-NLV-bulk-responsive donor “Gold” clones increased ~2-fold on average, while “Gold” clones from exog-NLV-bulk-non-responsive donors increased ~30-fold (**Figure 5D**). These data show that exog-NLV-bulk-responsive donors responded to CD3 co-potentiation by amplifying potent NLV-focused clones, while exog-NLV-bulk-non-responsive donors responded to combination treatment with synergy driven mostly by CD3 co-potentiation and low-but-positive intrinsic potency toward exogenous NLV. We conclude that Mono-OKT3-Fab provides antigen-specific CD3 co-potentiation that can increase expansion of recall public and private clones against antigens that are immunodominant or antigens of intrinsically weak potency (**Figure 6**).

**Figure 5.**
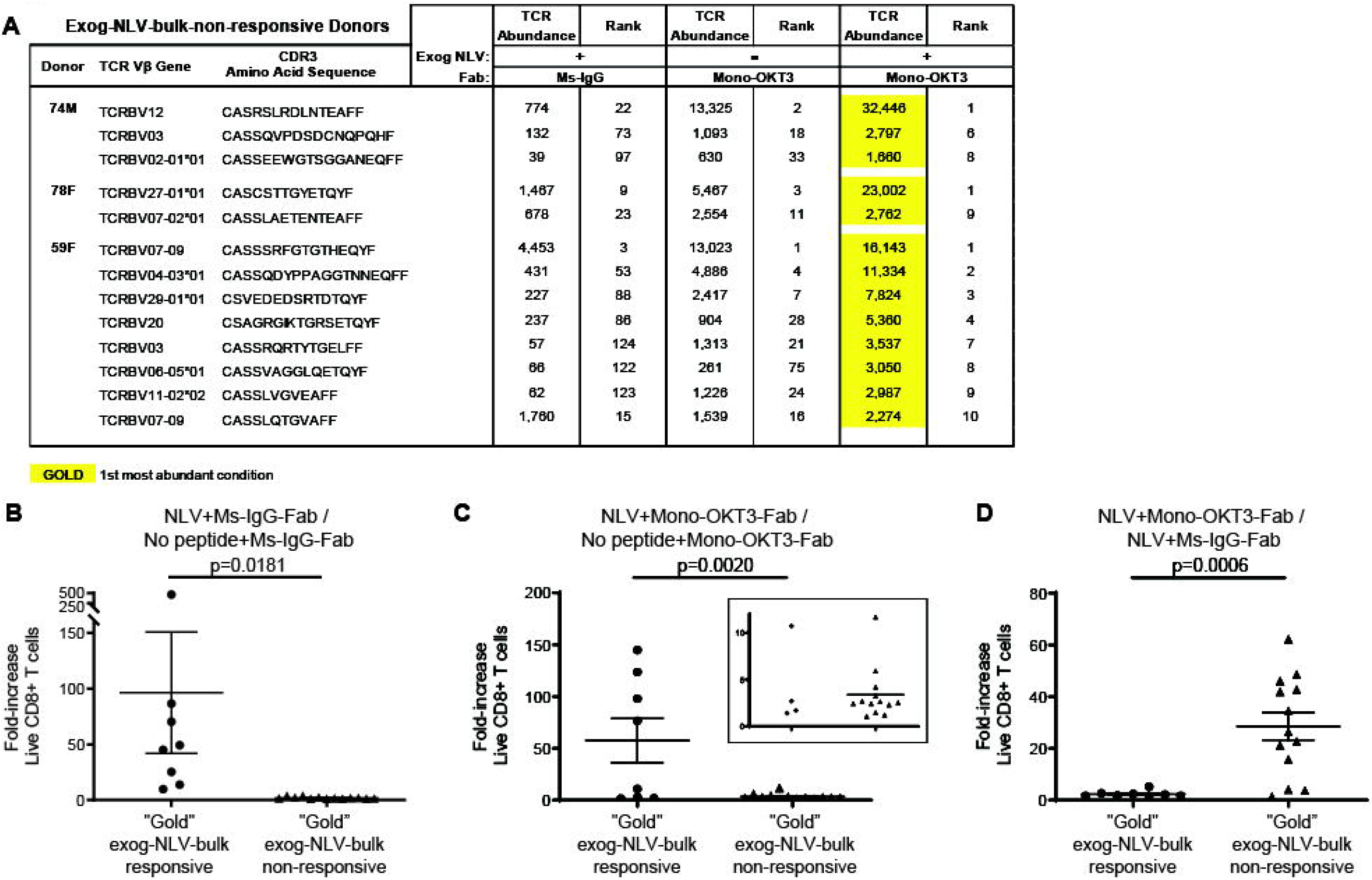
Although bulk cultures from A2/NLV-tetramer(-) donors appear non-responsive to exogenous NLV, single-clone analysis reveals weak NLV-responsive clones amplified by CD3 copotentiation in a unique expansion signature. (**A**) Gold/Silver-Bronze analysis was applied to exog-NLV-bulk-non-responsive donors for the top clones showing greater abundance in NLV+Ms-IgG-Fab versus No peptide+Ms-IgG-Fab conditions. It was observed that for each of these clones, the NLV+Mono-OKT3-Fab combination condition yielded greatest abundance. (**B**) Among top NLV-specific clones, those from exog-NLV-bulk-responsive donors respond more than those from exog-NLV-bulk-non-responsive donors to exogenous NLV. NLV-specific fold-increase in TCR abundance was determined for “Gold” responders from exog-NLV-bulk-responsive donors versus those from -non-responsive donors (Supplemental Table 5). (**C**) NLV-specific fold-increase in TCR abundance was also assessed when “Gold” response clones from both types of donors were cultured in the presence of Mono-OKT3-Fab. (**D**) Exog-NLV-bulk-non-responsive donors respond more than exog-NLV-bulk-responsive donors to CD3 co-potentiation when it is driven by exogenous NLV. Mono-OKT3-Fab-specific fold-increase in TCR abundance was determined for “Gold” responders from exog-NLV-bulk-responsive donors versus those from –non-responsive donors (Supplemental Table 5). Data are included for “Gold” clones in exog-NLV-bulk-responsive and -non-responsive donors. For (B-D), each dot represents the fold-increase of a TRBV-CDR3-bearing clone (mean ± SEM, one-tailed unpaired Student’s t-test).

## Discussion

We show that Mono-OKT3-Fab provides human CD3 co-potentiation to enhance expansion of several classes of recall CD8 T cells with relevance to HCMV. First, Mono-OKT3-Fab fulfilled the biochemical requirements to deliver co-potentiation: binding to CD3 and inducing CD3Δc without initiating intrinsic signaling or interfering with TCR:antigen binding (**Figures 1–2**). Functional copotentiation was observed in recall assays where PBMCs from healthy blood donors were cultured +/− exogenous NLV and/or Mono-OKT3-Fab. Because enhanced expansion was observed in both A2/NLV-tetramer(+) and A2/NLV-tetramer(-) cells, and from exog-NLV-bulk-non-responsive donors (**Figure 3A-C**), the question arose whether Mono-OKT3-Fab induced pan-T cell activation versus HLA-restricted copotentiation. Augmentation of expansion was impaired when using Mono-UCHTI-Fab, which binds CD3 (**Figure 1A**) and induces CD3Δc (**Figure 2C**), but inhibits TCR:antigen binding and signaling (**Figure 1B-C**), suggesting that Fab:CD3 was insufficient for co-potentiation without TCR:antigen interaction. Mono-OKT3-Fab-mediated co-potentiation was inhibited in the presence of anti-CD8 blocking antibody, suggesting that co-potentiation depends on the tripartite CD8:TCR:HLA antigenic interaction (**Figure 3D-F**). Furthermore, a pan-T cell stimulatory effect was not supported by clonal analysis, where, despite thousands of clones that were identified and measured, both co-potentiation and exogenous-peptide effects were mainly concentrated in top clones. Together, these results support a model in which Mono-OKT3-Fab performs co-potentiation by enhancing HLA-dependent responses (**Figure 6**). We speculate that autologous APCs present some antigens for which recall T cell clones are specific, and these are copotentiated by Mono-OKT3-Fab, while other recall T cells do not encounter their specific antigens on PBMC APCs, and these are not affected by Mono-OKT3-Fab.

**Figure 6.**
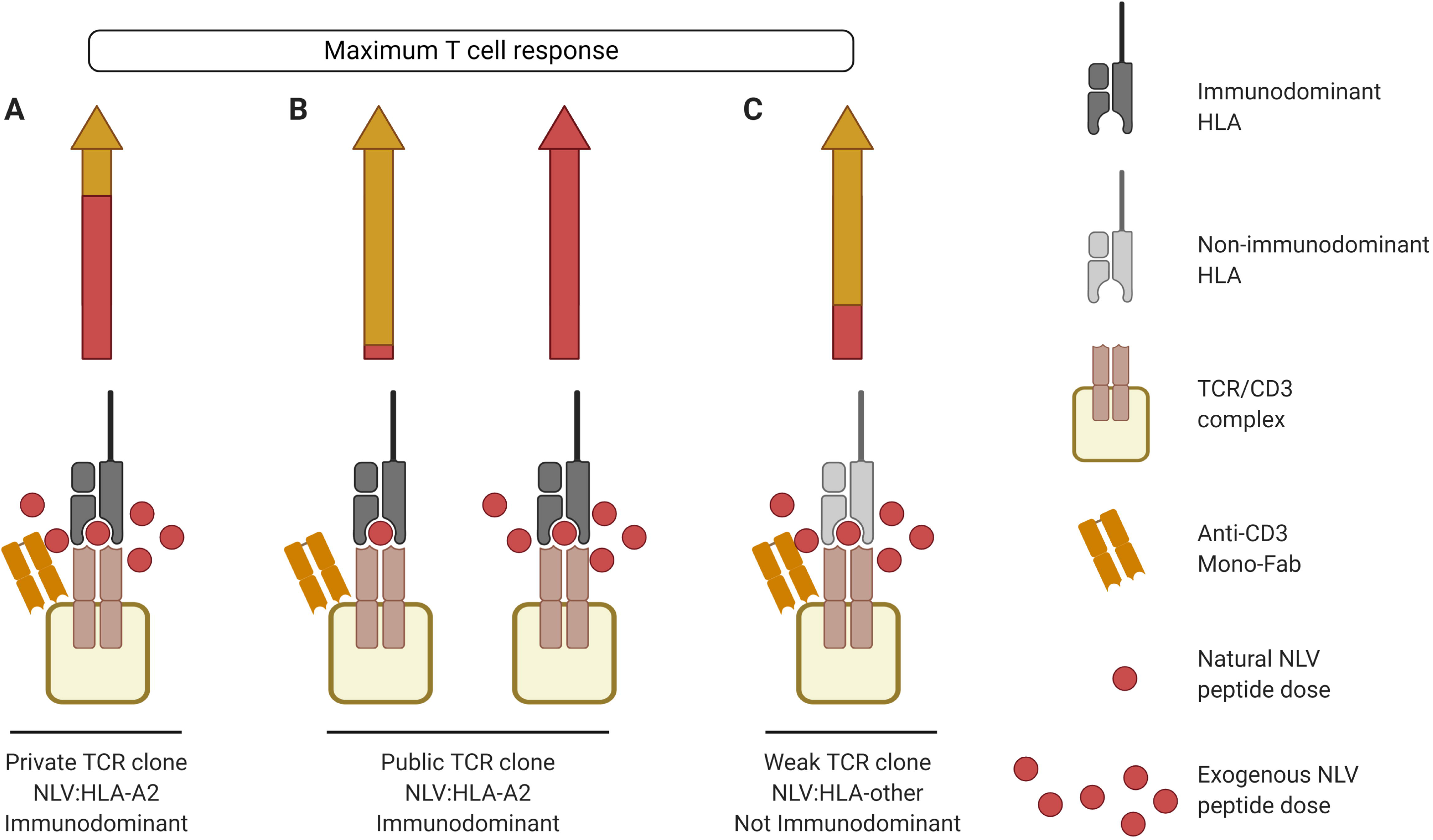
Different T cell clonal signatures of maximal recall response to NLV when providing copotentiation with anti-CD3 Mono-Fab. (**A**) Maximum recall response of private immunodominant TCR clones to exogenous NLV is mainly caused by the peptide (arrow, red segment), with a smaller contribution coming from co-potentiation delivered by anti-CD3 Mono-Fab (arrow, yellow segment). (**B**) Maximum recall immunodominant response of public TCR clones to NLV is driven by either (i) copotentiation (left arrow, yellow segment), with the smallest contribution from natural amounts of NLV presented in HCMV(+) APCs, or (ii) exogenous NLV alone (right arrow). (**C**) NLV weak TCR clones reach their maximum recall response to exogenous NLV mainly by co-potentiation (arrow, yellow segment), with a smaller contribution coming from exogenous NLV peptide (arrow, red segment). Created with BioRender.com.

Previous mouse experiments showed the CD3 co-potentiation principle in naïve CD8 T cells *in vitro,* and anti-tumor effects *in vivo*, without ruling out the possibility that other T cell classes and/or activation states could be responsive^11^. In the current work, studying recall CD8 T cells in response to NLV and Mono-OKT3-Fab, we observed co-potentiation of peripheral blood T cells in classic recall assays, suggesting that previously clonally expanded T cells are responsive to co-potentiation (**Figure 3**). Among them we observed public and private clones responding to immunodominant NLV:HLA-A2 antigen (**Figure 4; Table 1**), and clones for which NLV was a weak antigen (**Figure 5**).

Interestingly, these three classes of clonal response showed distinct clonal expansion signatures. A private signature against immunodominant NLV:HLA-A2 saw highest clonal abundance achieved upon combination stimulus of exogenous NLV+Mono-OKT3-Fab (**Table 1**). This was also observed for exog-NLV-bulk-non-responsive donors for whom NLV was an intrinsically weak antigen (**Figure 5**). The difference between these two response classes was in their opposite intrinsic potencies of NLV peptide (strong versus weak), and in a corresponding switch in contribution to the combinatorial synergistic response, peptide>Mono-Fab for immunodominance versus Mono-Fab>peptide for weak antigens (**Figure 5**). These patterns raise important questions. First, these responses can be confirmed as recall responses (not naïve), because negative-control culture conditions (and all culture conditions) contained multiple copies of these clones, consistent with prior clonal expansion. This is not surprising for responders to immunodominant antigen, but when NLV is a weak antigen, why would recall assays contain previously expanded T cell clones for which intrinsic reactivity to exogenous NLV is minimal? Three non-exclusive possibilities are that (i) over time *in vivo*, the weak response slowly accumulates numerous cells; (ii) *in vivo* the response to NLV is potent, although in recall assays it is not; (iii) clones were expanded *in vivo* against other antigens but they cross-react with NLV:HLA. This latter possibility is attractive because it has been suggested that heteroclitic stimulation of T cell clones might increase response to other cross-reactive antigens^36^, which could recruit pre-existing recall cells for other antigens to contribute to this otherwise non-sterilizing immune response.

The third expansion signature saw public T cells generating highest clonal abundance with either exogenous NLV or Mono-OKT3-Fab, but not with both in combination. The rest of the dataset reinforced the idea that the two modes of TCR/CD3 engagement were sterically and functionally compatible and synergistic, but apparently not in this case. One possible explanation is that the two modes of engagement remain synergistic, but they can cause hyperstimulation that favors activation-induced cell death^37,38^ or an anti-proliferative signal^39,40^, both of which involve TCR signals that are poorly understood, and both of which decrease clonal expansion. Why these responses might be favored in public more than private T cell clones is not clear, but leads to proposal of a second possible explanation. Perhaps public T cells possess a physical/functional distinction that has not been previously predicted, and their TCR/CD3 complexes engage or signal in response to cognate antigen and Mono-OKT3-Fab differently than most private TCR-clones. The nature of such a distinct property of the public T cell or its TCR/CD3 complex cannot be guessed at this time, but for now we report that public T cells responded with a unique clonal expansion pattern to CD3 co-potentiation.

Finally, the question of whether CD3 co-potentiation may be beneficial to patients merits further exploration. The present study provides compelling evidence that anti-human-CD3 Fabs can enhance expansion of several classes of recall T cell clones responding to antigens. It is conceivable that these effects may find useful translation *in vitro* for adoptive cell therapies, *in vivo* for immunoboosting, or in other immunotherapeutic strategies against chronic/persistent antigens in cancer or infectious diseases like HCMV.

## Supporting information

Graphical abstract

## Acknowledgements

We thank Balbino Alarcón (Universidad Autónoma de Madrid, Madrid, Spain) and Ed Palmer (University of Basel, Switzerland) for kind gifts of reagents. This work was funded by Mayo Graduate School of Biomedical Sciences (LREB), Mayo Foundation (SNM), NCI (U01 CA244314, DG; R33CA228979, AGS), NIAID (R01AI097187, DG), and NIGMS (R01GM103841, AGS).

## Authorship Contributions

LREB designed, performed, and analyzed experiments and wrote the paper. WKN, SSS, MA, MMH, CAP, and LRP designed, performed, analyzed, and/or interpreted data, and revised the paper. AGS and DG designed experiments, analyzed data, and wrote the paper. SNM designed experiments, analyzed and interpreted data, wrote the paper, and conceived the project.

