## Supplemental Tables and Figures for "Public and private human T cell clones respond differentially to HCMV antigen when boosted by CD3 co-potentiation"

| Donor | HLA-A*02:01 | A2/NLV-tetramer | Category |
|---|---|---|---|
| **72F** | **+** | **+** | **Exog-NLV-bulk-responsive** |
| **53M** | **+** | **+** | **Exog-NLV-bulk-responsive** |
| **28M** | **+** | **+** | **Exog-NLV-bulk-responsive** |
| **47M** | **+** | **+** | **Exog-NLV-bulk-responsive** |
| **74M** | **+** | **-** | **Exog-NLV-bulk-non-responsive** |
| **78F** | **-** | **-** | **Exog-NLV-bulk-non-responsive** |
| **59F** | **-** | **-** | **Exog-NLV-bulk-non-responsive** |

**Supplemental Table 1. PBMC Donors.** PBMCs from healthy donors were screened for HLA-A*02:01 positivity and for the presence of CD8 T cells specific for the HCMV pp65 peptide, NLVPMVATV (NLV), by tetramer staining. Each donor is distinguished by sex and age (years) at the time of blood draw. Categorization is based on whether recall CD8 T cell numbers increased in bulk culture in response to exogenous NLV relative to parallel cultures with no peptide added.

### Top-10 TCR analysis and commentary

1. Genomic DNA was sequenced on the last day (day 9) of recall assays of PBMCs from each donor, having been cultured +/- exogenous NLV peptide and control Ms-IgG-Fab or Mono-OKT3-Fab.

2. The frequencies of individual TCR clone sequences along with live cell counts from flow cytometry were used to determine the number of each unique TCR-bearing clone (TCR abundance) at day nine for each sample (see figure to the right).

3. TCR clones with the same TCR abundance were equally ranked. (For example, TCR abundances of 9000, 7500, 7500, 3000 would be ranked 1, 2, 2, 3, respectively). Importantly, all top ranks had only one clone, whereas bottom ranks had many clones sharing low copy number. At the top of each spreadsheet, "Rank of X" shows how many total ranks there were for each particular condition. "Rank percentile" was calculated by taking the TCR clone rank divided by the total ranks for that condition.

4. For each donor, TCR clone sequences were ranked by most abundant to least abundant in each of the four culture conditions, which are listed at the top of each sheet. The condition that was ranked most to least abundant is highlighted in yellow, and the TCR abundance and rank of that same amino acid sequence in the other three conditions are also shown. Top 10 ranks for each condition are included in this supplemental table.

5. TCR clones for which the NLV+Ms-IgG-Fab condition > No peptide+Ms-IgG-Fab condition by at least 10% were considered NLV-responsive, or for public TCRs, observation of that pattern in at least one other donor or were previously reported (highlighted in blue).

6. Exogenous-NLV-non-responsive clones are in red font. Some TCRs had high relative abundance in the No peptide+Ms-IgG-Fab negative-control condition, and exogenous NLV did not increase cell numbers. These clones are not considered exogenous-NLV-responsive.

**7. Public TCRs are bolded, defined as TCR clones that were either present in more than one donor, or else previously reported.**

8. Mint green highlighting denotes TCR sequences that are not considered exogenous-NLV-responsive (TCR abundance of NLV+Ms-IgG-Fab was not greater than the abundance of that clone in No peptide+Ms-IgG-Fab) even though the sequence appeared in the Top 10 ranks of the NLV+Ms-IgG-Fab condition.

9. Some top clones could not be accurately assessed for NLV-responsiveness due to failure to sample the same clones in control or comparator conditions. These clones, with a TCR abundance of 0 in one of the four conditions, are in black font without highlighting. However, if the TCR was a public TCR and showed NLV-responsiveness in at least one other donor or was previously published, then the clone was considered NLV-responsive.

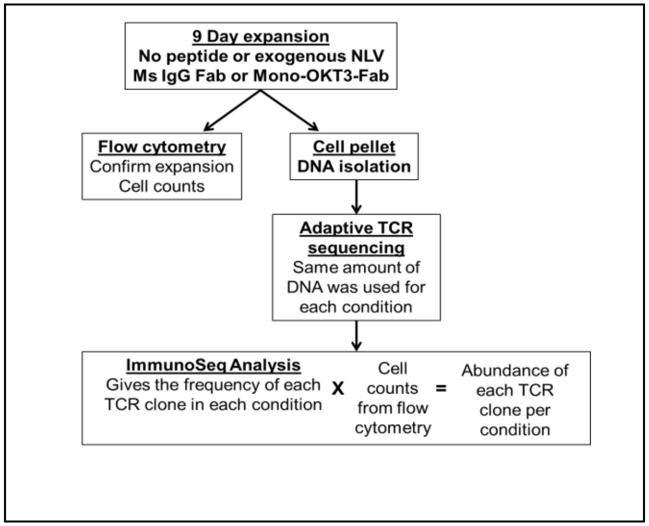


### Supplemental Table 2. TCR abundance and clone ranks from Exog-NLV-bulk-responsive donors.

| Exog-NLV-bulk- responsive donor 72F | | No peptide+Ms-IgG-Fab | | | NLV+Ms-IgG-Fab | | | No peptide+Mono-OKT3-Fab | | | NLV+Mono-OKT3-Fab | | |
|---|---|---|---|---|---|---|---|---|---|---|---|---|---|
| **TCR V beta gene** | **CDR3 Amino Acid Sequence** | **TCR Abundance** | **Rank of 142** | **Rank Percentile** | **TCR Abundance** | **Rank of 116** | **Rank Percentile** | **TCR Abundance** | **Rank of 142** | **Rank Percentile** | **TCR Abundance** | **Rank of 65** | **Rank Percentile** |
| TCRBV30-01*01 | CAWRPASYEQYF | 8,825 | 1 | 1% | 0 | 116 | 100% | 0 | 142 | 100% | 0 | 65 | 100% |
| TCRBV02-01*01 | CASILEDGGRNSPLHF | 5,804 | 2 | 1% | 0 | 116 | 100% | 0 | 142 | 100% | 0 | 65 | 100% |
| TCRBV07-09 | CASSPRNQGYEQYF | 4,988 | 3 | 2% | 0 | 116 | 100% | 1,141 | 26 | 18% | 247 | 40 | 62% |
| TCRBV12 | CASRDRENTEAFF | 4,796 | 4 | 3% | 2,280 | 8 | 7% | 3,583 | 7 | 5% | 3,871 | 8 | 12% |
| TCRBV07-09 | CASSLEQAYGYTF | 4,710 | 5 | 4% | 0 | 116 | 100% | 0 | 142 | 100% | 0 | 65 | 100% |
| TCRBV06 | CASSRSRDRDLYEQYF | 4,632 | 6 | 4% | 0 | 116 | 100% | 0 | 142 | 100% | 0 | 65 | 100% |
| TCRBV03 | CASSHRGSNQPQHF | 4,507 | 7 | 5% | 0 | 116 | 100% | 0 | 142 | 100% | 0 | 65 | 100% |
| TCRBV04-02*01 | CASSQDYSNQPQHF | 4,215 | 8 | 6% | 0 | 116 | 100% | 0 | 142 | 100% | 0 | 65 | 100% |
| TCRBV18-01*01 | CASSLSPDSYEQYF | 4,097 | 9 | 6% | 0 | 116 | 100% | 0 | 142 | 100% | 0 | 65 | 100% |
| TCRBV20 | CSARDPQVREQYF | 4,069 | 10 | 7% | 0 | 116 | 100% | 0 | 142 | 100% | 0 | 65 | 100% |
| **TCR V beta gene** | **CDR3 Amino Acid Sequence** | **TCR Abundance** | **Rank of 142** | **Rank Percentile** | **TCR Abundance** | **Rank of 116** | **Rank Percentile** | **TCR Abundance** | **Rank of 142** | **Rank Percentile** | **TCR Abundance** | **Rank of 65** | **Rank Percentile** |
| TCRBV06 | CASSKVTGTGNYGYTF | 1,511 | 26 | 18% | 130,547 | 1 | 1% | 125,632 | 1 | 1% | 214,687 | 1 | 2% |
| TCRBV06-05*01 | CASSYRGLTGELFF | 0 | 142 | 100% | 10,024 | 2 | 2% | 3,552 | 8 | 6% | 317 | 35 | 54% |
| TCRBV06-05*01 | CASSPSTGTIYGYTF | 118 | 109 | 77% | 8,254 | 3 | 3% | 7,241 | 4 | 3% | 19,561 | 3 | 5% |
| TCRBV06-05*01 | CASSYVGTGGGSAFF | 0 | 142 | 100% | 4,625 | 4 | 3% | 2,720 | 12 | 8% | 0 | 65 | 100% |
| TCRBV06 | CASSPITGQGAYGYTF | 428 | 60 | 42% | 4,163 | 5 | 4% | 6,263 | 5 | 4% | 8,800 | 4 | 6% |
| TCRBV05-01*01 | CASRFQPNTEAFF | 0 | 142 | 100% | 3,904 | 6 | 5% | 2,994 | 11 | 8% | 0 | 65 | 100% |
| TCRBV12 | CASSRDRGLGPQHF | 0 | 142 | 100% | 3,503 | 7 | 6% | 3,145 | 9 | 6% | 0 | 65 | 100% |
| TCRBV12 | CASRDRENTEAFF | 4,796 | 4 | 3% | 2,280 | 8 | 7% | 3,583 | 7 | 5% | 3,871 | 8 | 12% |
| TCRBV07-09 | CAGVYRVSNQPQHF | 0 | 142 | 100% | 2,114 | 9 | 8% | 1,455 | 19 | 13% | 0 | 65 | 100% |
| TCRBV12 | CASSLFRSAGNYGYTF | 0 | 142 | 100% | 1,665 | 10 | 9% | 1,672 | 16 | 11% | 0 | 65 | 100% |
| **TCR V beta gene** | **CDR3 Amino Acid Sequence** | **TCR Abundance** | **Rank of 142** | **Rank Percentile** | **TCR Abundance** | **Rank of 116** | **Rank Percentile** | **TCR Abundance** | **Rank of 142** | **Rank Percentile** | **TCR Abundance** | **Rank of 65** | **Rank Percentile** |
| TCRBV06 | CASSKVTGTGNYGYTF | 1,511 | 26 | 18% | 130,547 | 1 | 1% | 125,632 | 1 | 1% | 214,687 | 1 | 2% |
| TCRBV12 | CASSLALNTEAFF | 3,926 | 11 | 8% | 1,219 | 17 | 15% | 22,513 | 2 | 1% | 30,985 | 2 | 3% |
| TCRBV06 | CATFEEPNEKLFF | 1,457 | 27 | 19% | 834 | 25 | 22% | 7,426 | 3 | 2% | 7,207 | 5 | 8% |
| TCRBV06-05*01 | CASSPSTGTIYGYTF | 118 | 109 | 77% | 8,254 | 3 | 3% | 7,241 | 4 | 3% | 19,561 | 3 | 5% |
| TCRBV06 | CASSPITGQGAYGYTF | 428 | 60 | 42% | 4,163 | 5 | 4% | 6,263 | 5 | 4% | 8,800 | 4 | 6% |
| TCRBV19-01 | CASSIDSPNTEAFF | 677 | 45 | 32% | 340 | 54 | 47% | 4,180 | 6 | 4% | 2,703 | 9 | 14% |
| TCRBV12 | CASRDRENTEAFF | 4,796 | 4 | 3% | 2,280 | 8 | 7% | 3,583 | 7 | 5% | 3,871 | 8 | 12% |
| TCRBV06-05*01 | CASSYRGLTGELFF | 0 | 142 | 100% | 10,024 | 2 | 2% | 3,552 | 8 | 6% | 317 | 35 | 54% |
| TCRBV12 | CASSRDRGLGPQHF | 0 | 142 | 100% | 3,503 | 7 | 6% | 3,145 | 9 | 6% | 0 | 65 | 100% |
| TCRBV06-05*01 | CASSYADRGAGELFF | 937 | 34 | 24% | 794 | 26 | 22% | 3,109 | 10 | 7% | 4,465 | 7 | 11% |
| **TCR V beta gene** | **CDR3 Amino Acid Sequence** | **TCR Abundance** | **Rank of 142** | **Rank Percentile** | **TCR Abundance** | **Rank of 116** | **Rank Percentile** | **TCR Abundance** | **Rank of 142** | **Rank Percentile** | **TCR Abundance** | **Rank of 65** | **Rank Percentile** |
| TCRBV06 | CASSKVTGTGNYGYTF | 1,511 | 26 | 18% | 130,547 | 1 | 1% | 125,632 | 1 | 1% | 214,687 | 1 | 2% |
| TCRBV12 | CASSLALNTEAFF | 3,926 | 11 | 8% | 1,219 | 17 | 15% | 22,513 | 2 | 1% | 30,985 | 2 | 3% |
| TCRBV06-05*01 | CASSPSTGTIYGYTF | 118 | 109 | 77% | 8,254 | 3 | 3% | 7,241 | 4 | 3% | 19,561 | 3 | 5% |
| TCRBV06 | CASSPITGQGAYGYTF | 428 | 60 | 42% | 4,163 | 5 | 4% | 6,263 | 5 | 4% | 8,800 | 4 | 6% |
| TCRBV06 | CATFEEPNEKLFF | 1,457 | 27 | 19% | 834 | 25 | 22% | 7,426 | 3 | 2% | 7,207 | 5 | 8% |
| TCRBV11-02*02 | CASSPWAYATDTQYF | 29 | 134 | 94% | 1,284 | 15 | 13% | 619 | 39 | 27% | 6,652 | 6 | 9% |
| TCRBV06-05*01 | CASSYADRGAGELFF | 937 | 34 | 24% | 794 | 26 | 22% | 3,109 | 10 | 7% | 4,465 | 7 | 11% |
| TCRBV12 | CASRDRENTEAFF | 4,796 | 4 | 3% | 2,280 | 8 | 7% | 3,583 | 7 | 5% | 3,871 | 8 | 12% |
| TCRBV19-01 | CASSIDSPNTEAFF | 677 | 45 | 32% | 340 | 54 | 47% | 4,180 | 6 | 4% | 2,703 | 9 | 14% |
| TCRBV02-01*01 | CARTGYEDTEAFF | 0 | 142 | 100% | 0 | 116 | 100% | 0 | 142 | 100% | 1,435 | 10 | 15% |

**Supplemental Table 2. TCR abundance and clone ranks from Exog-NLV-bulk-responsive donors.**

| Exog-NLV-bulk- responsive donor 53M | | No peptide+Ms-IgG-Fab | | | NLV+Ms-IgG-Fab | | | No peptide+Mono-OKT3-Fab | | | NLV+Mono-OKT3-Fab | | | |
|---|---|---|---|---|---|---|---|---|---|---|---|---|---|---|
| **TCR V beta gene** | **CDR3 Amino Acid Sequence** | **TCR Abundance** | **Rank of 173** | **Rank Percentile** | **TCR Abundance** | **Rank of 139** | **Rank Percentile** | **TCR Abundance** | **Rank of 129** | **Rank Percentile** | **TCR Abundance** | **Rank of 100** | **Rank Percentile** | |
| TCRBV12 | CASSDLAGGSYEQYF | 21,528 | 1 | 1% | 0 | 139 | 100% | 10 | 128 | 99% | 0 | 100 | 100% | |
| TCRBV19-01 | CAIFLPGQGRDTQYF | 17,117 | 2 | 1% | 8,047 | 10 | 7% | 1,235 | 38 | 29% | 1,879 | 23 | 23% | |
| TCRBV29-01*01 | CSVEGGERVQETQYF | 16,446 | 3 | 2% | 8,706 | 8 | 6% | 1,842 | 28 | 22% | 2,515 | 16 | 16% | |
| TCRBV12 | CASAGETGELFF | 12,443 | 4 | 2% | 5,785 | 15 | 11% | 1,564 | 33 | 26% | 737 | 51 | 51% | |
| TCRBV05-01*01 | CASSLYGMNTEAFF | 10,233 | 5 | 3% | 0 | 139 | 100% | 0 | 129 | 100% | 0 | 100 | 100% | |
| TCRBV04-01*01 | CASSQDTGDEQYF | 7,012 | 6 | 3% | 23 | 137 | 99% | 10 | 128 | 99% | 0 | 100 | 100% | |
| TCRBV12 | CASSWDRNYGYTF | 6,689 | 7 | 4% | 11 | 138 | 99% | 20 | 127 | 98% | 0 | 100 | 100% | |
| TCRBV05-08*01 | CASSRKGEAFF | 5,558 | 8 | 5% | 3,080 | 27 | 19% | 50 | 124 | 96% | 101 | 93 | 93% | |
| TCRBV03 | CASSQDSRGLSYNEQFF | 5,303 | 9 | 5% | 0 | 139 | 100% | 10 | 128 | 99% | 0 | 100 | 100% | |
| TCRBV07-06*01 | CASSSGGTYGYTF | 4,955 | 10 | 6% | 796 | 74 | 53% | 50 | 124 | 96% | 43 | 97 | 97% | |
| **TCR V beta gene** | **CDR3 Amino Acid Sequence** | **TCR Abundance** | **Rank of 173** | **Rank Percentile** | **TCR Abundance** | **Rank of 139** | **Rank Percentile** | **TCR Abundance** | **Rank of 129** | **Rank Percentile** | **TCR Abundance** | **Rank of 100** | **Rank Percentile** | |
| TCRBV05-08*01 | CASSRTSINEQFF | 110 | 160 | 92% | 52,021 | 1 | 1% | 627 | 67 | 52% | 90,986 | 1 | 1% | |
| **TCRBV06** | **CASSPTTGTGTYGYTF** | 25 | 170 | 98% | 19,163 | 2 | 1% | 42,683 | 1 | 1% | 16,740 | 3 | 3% | |
| TCRBV28-01*01 | CASSPPGQSYEQYF | 0 | 173 | 100% | 15,435 | 3 | 2% | 0 | 129 | 100% | 0 | 100 | 100% | |
| TCRBV07-09 | CASSLGEGTTKAFF | 0 | 173 | 100% | 13,696 | 4 | 3% | 0 | 129 | 100% | 0 | 100 | 100% | |
| TCRBV07-09 | CASSLGGLDYGYTF | 0 | 173 | 100% | 12,616 | 5 | 4% | 0 | 129 | 100% | 0 | 100 | 100% | |
| **TCRBV12** | **CASSPGTYGYTF** | 0 | 173 | 100% | 10,945 | 6 | 4% | 458 | 84 | 65% | 6,158 | 6 | 6% | *** |
| **TCRBV27-01*01** | **CASSPIAGYPHEQYF** | 0 | 173 | 100% | 10,604 | 7 | 5% | 15,904 | 4 | 3% | 8,197 | 5 | 5% | *** |
| TCRBV29-01*01 | CSVEGGERVQETQYF | 16,446 | 3 | 2% | 8,706 | 8 | 6% | 1,842 | 28 | 22% | 2,515 | 16 | 16% | |
| TCRBV28-01*01 | CASSPPGQRYEQYF | 0 | 173 | 100% | 8,695 | 9 | 6% | 0 | 129 | 100% | 0 | 100 | 100% | |
| TCRBV19-01 | CAIFLPGQGRDTQYF | 17,117 | 2 | 1% | 8,047 | 10 | 7% | 1,235 | 38 | 29% | 1,879 | 23 | 23% | |
| **TCR V beta gene** | **CDR3 Amino Acid Sequence** | **TCR Abundance** | **Rank of 173** | **Rank Percentile** | **TCR Abundance** | **Rank of 139** | **Rank Percentile** | **TCR Abundance** | **Rank of 129** | **Rank Percentile** | **TCR Abundance** | **Rank of 100** | **Rank Percentile** | |
| **TCRBV06** | **CASSPTTGTGTYGYTF** | 25 | 170 | 98% | 19,163 | 2 | 1% | 42,683 | 1 | 1% | 16,740 | 3 | 3% | |
| **TCRBV12** | **CASSIMNEQFF** | 0 | 173 | 100% | 7,183 | 12 | 9% | 21,212 | 2 | 2% | 5,276 | 7 | 7% | *** |
| TCRBV04-01*01 | CASSPEGGGGAFF | 1,249 | 62 | 36% | 852 | 72 | 52% | 16,282 | 3 | 2% | 19,386 | 2 | 2% | |
| **TCRBV27-01*01** | **CASSPIAGYPHEQYF** | 0 | 173 | 100% | 10,604 | 7 | 5% | 15,904 | 4 | 3% | 8,197 | 5 | 5% | *** |
| **TCRBV12** | **CASSSVNEQFF** | 8 | 172 | 99% | 3,535 | 22 | 16% | 11,423 | 5 | 4% | 3,050 | 15 | 15% | |
| **TCRBV12** | **CASSSAYYGYTF** | 8 | 172 | 99% | 3,148 | 25 | 18% | 9,461 | 6 | 5% | 3,123 | 14 | 14% | |
| **TCRBV12** | **CASSSVNEQYF** | 0 | 173 | 100% | 2,307 | 31 | 22% | 6,473 | 7 | 5% | 1,980 | 21 | 21% | *** |
| TCRBV28-01*01 | CASHNPSLSYEQYF | 0 | 173 | 100% | 0 | 139 | 100% | 5,686 | 8 | 6% | 0 | 100 | 100% | |
| TCRBV11-01*01 | CASSLAPPYEQYF | 195 | 150 | 87% | 148 | 126 | 91% | 5,527 | 9 | 7% | 4,742 | 8 | 8% | |
| TCRBV20 | CSAVRPPNQPQHF | 178 | 152 | 88% | 341 | 109 | 78% | 4,740 | 10 | 8% | 2,125 | 19 | 19% | |
| **TCR V beta gene** | **CDR3 Amino Acid Sequence** | **TCR Abundance** | **Rank of 173** | **Rank Percentile** | **TCR Abundance** | **Rank of 139** | **Rank Percentile** | **TCR Abundance** | **Rank of 129** | **Rank Percentile** | **TCR Abundance** | **Rank of 100** | **Rank Percentile** | |
| TCRBV05-08*01 | CASSRTSINEQFF | 110 | 160 | 92% | 52,021 | 1 | 1% | 627 | 67 | 52% | 90,986 | 1 | 1% | |
| TCRBV04-01*01 | CASSPEGGGGAFF | 1,249 | 62 | 36% | 852 | 72 | 52% | 16,282 | 3 | 2% | 19,386 | 2 | 2% | |
| **TCRBV06** | **CASSPTTGTGTYGYTF** | 25 | 170 | 98% | 19,163 | 2 | 1% | 42,683 | 1 | 1% | 16,740 | 3 | 3% | |
| **TCRBV27-01*01** | **CASSLEGYTEAFF** | 340 | 133 | 77% | 4,717 | 18 | 13% | 100 | 119 | 92% | 12,317 | 4 | 4% | |
| **TCRBV27-01*01** | **CASSPIAGYPHEQYF** | 0 | 173 | 100% | 10,604 | 7 | 5% | 15,904 | 4 | 3% | 8,197 | 5 | 5% | *** |
| **TCRBV12** | **CASSPGTYGYTF** | 0 | 173 | 100% | 10,945 | 6 | 4% | 458 | 84 | 65% | 6,158 | 6 | 6% | *** |
| **TCRBV12** | **CASSIMNEQFF** | 0 | 173 | 100% | 7,183 | 12 | 9% | 21,212 | 2 | 2% | 5,276 | 7 | 7% | *** |
| TCRBV11-01*01 | CASSLAPPYEQYF | 195 | 150 | 87% | 148 | 126 | 91% | 5,527 | 9 | 7% | 4,742 | 8 | 8% | |
| TCRBV04-01*01 | CASSQGVGLGEKLFF | 484 | 116 | 67% | 318 | 111 | 80% | 3,426 | 17 | 13% | 4,192 | 9 | 9% | |
| TCRBV09-01 | CASSPRDNPNYGYTF | 0 | 173 | 100% | 0 | 139 | 100% | 0 | 129 | 100% | 3,831 | 10 | 10% | |

***** These public TCR clones are considered NLV-responsive because they were observed as exog-NLV-responsive in at least one other donor, or previously reported in the literature**

### Supplemental Table 2. TCR abundance and clone ranks from Exog-NLV-bulk-responsive donors.

| Exog-NLV-bulk- responsive donor 28M | | No peptide+Ms-IgG-Fab | | | NLV+Ms-IgG-Fab | | | No peptide+Mono-OKT3-Fab | | | NLV+Mono-OKT3-Fab | | |
|---|---|---|---|---|---|---|---|---|---|---|---|---|---|
| TCR V beta gene | CDR3 Amino Acid Sequence | TCR Abundance | Rank of 102 | Rank Percentile | TCR Abundance | Rank of 92 | Rank Percentile | TCR Abundance | Rank of 123 | Rank Percentile | TCR Abundance | Rank of 109 | Rank Percentile |
| TCRBV28-01*01 | CASSPPGQARAIYGYTF | 10,873 | 1 | 1% | 496 | 36 | 39% | 0 | 123 | 100% | 0 | 109 | 100% |
| TCRBV20 | CSARDKASGMNSYNEQFF | 5,250 | 2 | 2% | 3,230 | 7 | 8% | 1,866 | 18 | 15% | 1,900 | 20 | 18% |
| TCRBV27-01*01 | CASRDGGYEQYF | 4,960 | 3 | 3% | 1,644 | 13 | 14% | 2,324 | 13 | 11% | 7,012 | 6 | 6% |
| TCRBV12 | CASSINYSNQPQHF | 3,984 | 4 | 4% | 1,943 | 12 | 13% | 3,944 | 7 | 6% | 4,909 | 10 | 9% |
| TCRBV29-01*01 | CSARDGSSNQPQHF | 3,055 | 5 | 5% | 699 | 28 | 30% | 294 | 62 | 50% | 977 | 32 | 29% |
| TCRBV12-05*01 | CASGLGVNTEAFF | 2,933 | 6 | 6% | 1,174 | 16 | 17% | 4,199 | 5 | 4% | 7,153 | 5 | 5% |
| TCRBV19-01 | CASSSRTSGRFYEQYF | 2,461 | 7 | 7% | 881 | 18 | 20% | 4,173 | 6 | 5% | 5,848 | 8 | 7% |
| TCRBV09-01 | CASSLRLWDTEAFF | 1,836 | 8 | 8% | 0 | 92 | 100% | 0 | 123 | 100% | 0 | 109 | 100% |
| TCRBV06 | CASGDTQYF | 1,681 | 9 | 9% | 0 | 92 | 100% | 0 | 123 | 100% | 0 | 109 | 100% |
| TCRBV04-01*01 | CASSQDLRTASYEQYF | 1,555 | 10 | 10% | 91 | 76 | 83% | 125 | 94 | 76% | 16 | 107 | 98% |
| TCR V beta gene | CDR3 Amino Acid Sequence | TCR Abundance | Rank of 102 | Rank Percentile | TCR Abundance | Rank of 92 | Rank Percentile | TCR Abundance | Rank of 123 | Rank Percentile | TCR Abundance | Rank of 109 | Rank Percentile |
| **TCRBV12** | **CASSSVNEQFF** | 1,224 | 12 | 12% | 90,590 | 1 | 1% | 10,008 | 2 | 2% | 57,318 | 1 | 1% |
| TCRBV06-05*01 | CASSPKTGATYGYTF | 518 | 26 | 25% | 22,100 | 2 | 2% | 2,829 | 12 | 10% | 22,061 | 2 | 2% |
| **TCRBV06** | **CASSPTTGTGTYGYTF** | 215 | 59 | 58% | 7,965 | 3 | 3% | 19,982 | 1 | 1% | 11,132 | 4 | 4% |
| TCRBV10-03*01 | CAISSPGSYGYTF | 0 | 102 | 100% | 7,618 | 4 | 4% | 0 | 123 | 100% | 0 | 109 | 100% |
| TCRBV06 | CASTPQTGTGYYGYTF | 294 | 48 | 47% | 7,420 | 5 | 5% | 160 | 86 | 70% | 12,250 | 3 | 3% |
| **TCRBV27-01*01** | **CASSPIAGYPHEQYF** | 47 | 92 | 90% | 3,779 | 6 | 7% | 7,646 | 3 | 2% | 6,004 | 7 | 6% |
| TCRBV20 | CSARDKASGMNSYNEQFF | 5,250 | 2 | 2% | 3,230 | 7 | 8% | 1,866 | 18 | 15% | 1,900 | 20 | 18% |
| TCRBV07-09 | CASSAGGSSYEQYF | 0 | 102 | 100% | 2,808 | 8 | 9% | 0 | 123 | 100% | 0 | 109 | 100% |
| TCRBV11-02*02 | CASSRRLAGDTGELFF | 710 | 18 | 18% | 2,755 | 9 | 10% | 2,155 | 16 | 13% | 297 | 71 | 65% |
| TCRBV29-01*01 | CSVAGGGYEQYF | 0 | 102 | 100% | 2,712 | 10 | 11% | 0 | 123 | 100% | 0 | 109 | 100% |
| TCR V beta gene | CDR3 Amino Acid Sequence | TCR Abundance | Rank of 102 | Rank Percentile | TCR Abundance | Rank of 92 | Rank Percentile | TCR Abundance | Rank of 123 | Rank Percentile | TCR Abundance | Rank of 109 | Rank Percentile |
| **TCRBV06** | **CASSPTTGTGTYGYTF** | 215 | 59 | 58% | 7,965 | 3 | 3% | 19,982 | 1 | 1% | 11,132 | 4 | 4% |
| **TCRBV12** | **CASSSVNEQFF** | 1,224 | 12 | 12% | 90,590 | 1 | 1% | 10,008 | 2 | 2% | 57,318 | 1 | 1% |
| **TCRBV27-01*01** | **CASSPIAGYPHEQYF** | 47 | 92 | 90% | 3,779 | 6 | 7% | 7,646 | 3 | 2% | 6,004 | 7 | 6% |
| **TCRBV12** | **CASSIMNEQFF** | 70 | 87 | 85% | 2,365 | 11 | 12% | 7,404 | 4 | 3% | 3,917 | 11 | 10% |
| TCRBV12-05*01 | CASGLGVNTEAFF | 2,933 | 6 | 6% | 1,174 | 16 | 17% | 4,199 | 5 | 4% | 7,153 | 5 | 5% |
| TCRBV19-01 | CASSSRTSGRFYEQYF | 2,461 | 7 | 7% | 881 | 18 | 20% | 4,173 | 6 | 5% | 5,848 | 8 | 7% |
| TCRBV12 | CASSINYSNQPQHF | 3,984 | 4 | 4% | 1,943 | 12 | 13% | 3,944 | 7 | 6% | 4,909 | 10 | 9% |
| TCRBV04-02*01 | CASSQGETSYEQYF | 472 | 29 | 28% | 133 | 68 | 74% | 3,516 | 8 | 7% | 3,604 | 12 | 11% |
| **TCRBV12** | **CASSSAYYGYTF** | 37 | 94 | 92% | 1,206 | 15 | 16% | 3,119 | 9 | 7% | 2,251 | 17 | 16% |
| TCRBV02-01*01 | CASSESPATSGIMGYEQYF | 1,102 | 15 | 15% | 336 | 43 | 47% | 3,084 | 10 | 8% | 3,596 | 13 | 12% |
| TCR V beta gene | CDR3 Amino Acid Sequence | TCR Abundance | Rank of 102 | Rank Percentile | TCR Abundance | Rank of 92 | Rank Percentile | TCR Abundance | Rank of 123 | Rank Percentile | TCR Abundance | Rank of 109 | Rank Percentile |
| **TCRBV12** | **CASSSVNEQFF** | 1,224 | 12 | 12% | 90,590 | 1 | 1% | 10,008 | 2 | 2% | 57,318 | 1 | 1% |
| TCRBV06-05*01 | CASSPKTGATYGYTF | 518 | 26 | 25% | 22,100 | 2 | 2% | 2,829 | 12 | 10% | 22,061 | 2 | 2% |
| TCRBV06 | CASTPQTGTGYYGYTF | 294 | 48 | 47% | 7,420 | 5 | 5% | 160 | 86 | 70% | 12,250 | 3 | 3% |
| **TCRBV06** | **CASSPTTGTGTYGYTF** | 215 | 59 | 58% | 7,965 | 3 | 3% | 19,982 | 1 | 1% | 11,132 | 4 | 4% |
| TCRBV12-05*01 | CASGLGVNTEAFF | 2,933 | 6 | 6% | 1,174 | 16 | 17% | 4,199 | 5 | 4% | 7,153 | 5 | 5% |
| TCRBV27-01*01 | CASRDGGYEQYF | 4,960 | 3 | 3% | 1,644 | 13 | 14% | 2,324 | 13 | 11% | 7,012 | 6 | 6% |
| **TCRBV27-01*01** | **CASSPIAGYPHEQYF** | 47 | 92 | 90% | 3,779 | 6 | 7% | 7,646 | 3 | 2% | 6,004 | 7 | 6% |
| TCRBV19-01 | CASSSRTSGRFYEQYF | 2,461 | 7 | 7% | 881 | 18 | 20% | 4,173 | 6 | 5% | 5,848 | 8 | 7% |
| TCRBV11-02*02 | CASSLDPSGRLGDEQYF | 107 | 79 | 77% | 69 | 79 | 86% | 3,063 | 11 | 9% | 5,582 | 9 | 8% |
| TCRBV12 | CASSINYSNQPQHF | 3,984 | 4 | 4% | 1,943 | 12 | 13% | 3,944 | 7 | 6% | 4,909 | 10 | 9% |

**Supplemental Table 2. TCR abundance and clone ranks from Exog-NLV-bulk-responsive donors.**

| Exog-NLV-bulk- responsive donor 47M | | No peptide+Ms-IgG-Fab | | | NLV+Ms-IgG-Fab | | | No peptide+Mono-OKT3-Fab | | | NLV+Mono-OKT3-Fab | | | Notes |
|---|---|---|---|---|---|---|---|---|---|---|---|---|---|---|
| **TCR V beta gene** | **CDR3 Amino Acid Sequence** | **TCR Abundance** | **Rank of 111** | **Rank Percentile** | **TCR Abundance** | **Rank of 115** | **Rank Percentile** | **TCR Abundance** | **Rank of 120** | **Rank Percentile** | **TCR Abundance** | **Rank of 110** | **Rank Percentile** | |
| **TCRBV14-01*01** | **CASSQEPGNYGYTF** | 7,087 | 1 | 1% | 4,922 | 6 | 5% | 20,268 | 4 | 3% | 18,324 | 7 | 6% | *** |
| **TCRBV29-01*01** | **CSVTLPQADGRYGYTF** | 5,202 | 2 | 2% | 4,512 | 7 | 6% | 44,657 | 2 | 2% | 40,088 | 3 | 3% | *** |
| **TCRBV20** | **CSANQGGGNTEAFF** | 4,615 | 3 | 3% | 3,636 | 8 | 7% | 54,735 | 1 | 1% | 40,691 | 2 | 2% | *** |
| TCRBV10-02*01 | CASSESGHGSYEQYF | 4,050 | 4 | 4% | 2,438 | 10 | 9% | 2,356 | 14 | 12% | 3,112 | 15 | 14% | |
| TCRBV04-01*01 | CASSQTRGAGNTIYF | 3,784 | 5 | 5% | 2,386 | 11 | 10% | 18,085 | 5 | 4% | 17,021 | 8 | 7% | |
| TCRBV06-05*01 | CASSLGQNTEAFF | 3,761 | 6 | 5% | 2,162 | 12 | 10% | 1,815 | 16 | 13% | 1,959 | 18 | 16% | |
| TCRBV20-01*01 | CSARPGQGAYF | 2,367 | 7 | 6% | 0 | 115 | 100% | 0 | 120 | 100% | 0 | 110 | 100% | |
| TCRBV07-02*01 | CASSYSLPGLYEQYF | 2,326 | 8 | 7% | 0 | 115 | 100% | 0 | 120 | 100% | 0 | 110 | 100% | |
| TCRBV05-06*01 | CASSLGTGLYGYTF | 2,248 | 9 | 8% | 1,331 | 15 | 13% | 542 | 44 | 37% | 963 | 29 | 26% | |
| TCRBV19-01 | CASRRQQEPYGYTF | 2,243 | 10 | 9% | 355 | 44 | 38% | 1,837 | 15 | 13% | 138 | 89 | 81% | |
| **TCR V beta gene** | **CDR3 Amino Acid Sequence** | **TCR Abundance** | **Rank of 111** | **Rank Percentile** | **TCR Abundance** | **Rank of 115** | **Rank Percentile** | **TCR Abundance** | **Rank of 120** | **Rank Percentile** | **TCR Abundance** | **Rank of 110** | **Rank Percentile** | |
| **TCRBV27-01*01** | **CASSPIAGYPHEQYF** | 913 | 19 | 17% | 51,372 | 1 | 1% | 4,238 | 10 | 8% | 34,592 | 4 | 4% | |
| **TCRBV06** | **CASSPTTGTGTYGYTF** | 638 | 31 | 28% | 31,414 | 2 | 2% | 497 | 47 | 39% | 48,671 | 1 | 1% | |
| **TCRBV12** | **CASSSAYYGYTF** | 128 | 83 | 75% | 23,084 | 3 | 3% | 341 | 63 | 53% | 21,567 | 5 | 5% | |
| **TCRBV12** | **CASSSVNEQFF** | 110 | 87 | 78% | 7,276 | 4 | 3% | 45 | 112 | 93% | 3,374 | 14 | 13% | |
| **TCRBV12** | **CASSIMNEQFF** | 197 | 68 | 61% | 6,722 | 5 | 4% | 184 | 87 | 73% | 3,708 | 12 | 11% | |
| **TCRBV14-01*01** | **CASSQEPGNYGYTF** | 7,087 | 1 | 1% | 4,922 | 6 | 5% | 20,268 | 4 | 3% | 18,324 | 7 | 6% | *** |
| **TCRBV29-01*01** | **CSVTLPQADGRYGYTF** | 5,202 | 2 | 2% | 4,512 | 7 | 6% | 44,657 | 2 | 2% | 40,088 | 3 | 3% | *** |
| **TCRBV20** | **CSANQGGGNTEAFF** | 4,615 | 3 | 3% | 3,636 | 8 | 7% | 54,735 | 1 | 1% | 40,691 | 2 | 2% | *** |
| **TCRBV12** | **CASSSVNEQYF** | 60 | 98 | 88% | 3,545 | 9 | 8% | 39 | 113 | 94% | 3,603 | 13 | 12% | |
| TCRBV10-02*01 | CASSESGHGSYEQYF | 4,050 | 4 | 4% | 2,438 | 10 | 9% | 2,356 | 14 | 12% | 3,112 | 15 | 14% | |
| **TCR V beta gene** | **CDR3 Amino Acid Sequence** | **TCR Abundance** | **Rank of 111** | **Rank Percentile** | **TCR Abundance** | **Rank of 115** | **Rank Percentile** | **TCR Abundance** | **Rank of 120** | **Rank Percentile** | **TCR Abundance** | **Rank of 110** | **Rank Percentile** | |
| **TCRBV20** | **CSANQGGGNTEAFF** | 4,615 | 3 | 3% | 3,636 | 8 | 7% | 54,735 | 1 | 1% | 40,691 | 2 | 2% | *** |
| **TCRBV29-01*01** | **CSVTLPQADGRYGYTF** | 5,202 | 2 | 2% | 4,512 | 7 | 6% | 44,657 | 2 | 2% | 40,088 | 3 | 3% | *** |
| TCRBV07-08*01 | CASSNPGGSSYYEQYF | 1,266 | 14 | 13% | 898 | 19 | 17% | 23,618 | 3 | 3% | 21,528 | 6 | 5% | |
| **TCRBV14-01*01** | **CASSQEPGNYGYTF** | 7,087 | 1 | 1% | 4,922 | 6 | 5% | 20,268 | 4 | 3% | 18,324 | 7 | 6% | *** |
| TCRBV04-01*01 | CASSQTRGAGNTIYF | 3,784 | 5 | 5% | 2,386 | 11 | 10% | 18,085 | 5 | 4% | 17,021 | 8 | 7% | |
| TCRBV07-08*01 | CASSLGQAYEQYF | 1,431 | 13 | 12% | 589 | 25 | 22% | 14,366 | 6 | 5% | 15,304 | 9 | 8% | |
| TCRBV06 | CASRPTGGASEAFF | 931 | 18 | 16% | 127 | 75 | 65% | 6,522 | 7 | 6% | 3,754 | 11 | 10% | |
| TCRBV27-01*01 | CASSFGPSHNEQFF | 385 | 46 | 41% | 212 | 60 | 52% | 5,098 | 8 | 7% | 2,811 | 16 | 15% | |
| TCRBV27-01*01 | CASRQGFPGNEQFF | 500 | 37 | 33% | 277 | 50 | 43% | 4,551 | 9 | 8% | 4,396 | 10 | 9% | |
| **TCRBV27-01*01** | **CASSPIAGYPHEQYF** | 913 | 19 | 17% | 51,372 | 1 | 1% | 4,238 | 10 | 8% | 34,592 | 4 | 4% | |
| **TCR V beta gene** | **CDR3 Amino Acid Sequence** | **TCR Abundance** | **Rank of 111** | **Rank Percentile** | **TCR Abundance** | **Rank of 115** | **Rank Percentile** | **TCR Abundance** | **Rank of 120** | **Rank Percentile** | **TCR Abundance** | **Rank of 110** | **Rank Percentile** | |
| **TCRBV06** | **CASSPTTGTGTYGYTF** | 638 | 31 | 28% | 31,414 | 2 | 2% | 497 | 47 | 39% | 48,671 | 1 | 1% | |
| **TCRBV20** | **CSANQGGGNTEAFF** | 4,615 | 3 | 3% | 3,636 | 8 | 7% | 54,735 | 1 | 1% | 40,691 | 2 | 2% | *** |
| **TCRBV29-01*01** | **CSVTLPQADGRYGYTF** | 5,202 | 2 | 2% | 4,512 | 7 | 6% | 44,657 | 2 | 2% | 40,088 | 3 | 3% | *** |
| **TCRBV27-01*01** | **CASSPIAGYPHEQYF** | 913 | 19 | 17% | 51,372 | 1 | 1% | 4,238 | 10 | 8% | 34,592 | 4 | 4% | |
| **TCRBV12** | **CASSSAYYGYTF** | 128 | 83 | 75% | 23,084 | 3 | 3% | 341 | 63 | 53% | 21,567 | 5 | 5% | |
| TCRBV07-08*01 | CASSNPGGSSYYEQYF | 1,266 | 14 | 13% | 898 | 19 | 17% | 23,618 | 3 | 3% | 21,528 | 6 | 5% | |
| **TCRBV14-01*01** | **CASSQEPGNYGYTF** | 7,087 | 1 | 1% | 4,922 | 6 | 5% | 20,268 | 4 | 3% | 18,324 | 7 | 6% | *** |
| TCRBV04-01*01 | CASSQTRGAGNTIYF | 3,784 | 5 | 5% | 2,386 | 11 | 10% | 18,085 | 5 | 4% | 17,021 | 8 | 7% | |
| TCRBV07-08*01 | CASSLGQAYEQYF | 1,431 | 13 | 12% | 589 | 25 | 22% | 14,366 | 6 | 5% | 15,304 | 9 | 8% | |
| TCRBV27-01*01 | CASRQGFPGNEQFF | 500 | 37 | 33% | 277 | 50 | 43% | 4,551 | 9 | 8% | 4,396 | 10 | 9% | |

***** These public TCR clones are considered NLV-responsive because they were observed as exog-NLV-responsive in at least one other donor, or previously reported in the literature**

### Supplemental Table 2. TCR abundance and clone ranks from Exog-NLV-bulk-responsive donors.

A

| TRBV gene | CDR3 Amino Acid Sequence | Donors 53M | 28M | 47M | Prior References |
|---|---|---|---|---|---|
| TCRBV06 | CASSPTTGTGTYGYTF | ✔ | ✔ | ✔ | |
| TCRBV14-01*01 | CASSQEPGNYGYTF | ✔ | ✔ | ✔ | |
| TCRBV20 | CSANQGGGNTEAFF | ✔ | ✔ | ✔ | |
| TCRBV29-01*01 | CSVTLPQADGRYGYTF | ✔ | ✔ | ✔ | |
| TCRBV27-01*01 | CASSPIAGYPHEQYF | ✔ | ✔ | ✔ | |
| TCRBV27-01*01 | CASSLEGYTEAFF | ✔ | | | 14, 18, 19, 24 |
| TCRBV12 | CASSPGTYGYTF | ✔ | | | 19, 24 |
| TCRBV12 | CASSIMNEQFF | ✔ | ✔ | ✔ | |
| TCRBV12 | CASSSAYYGYTF | ✔ | ✔ | ✔ | 14, 19, 23, 24, 41 |
| TCRBV12 | CASSSVNEQFF | ✔ | ✔ | ✔ | 17, 22, 24, 41 |
| TCRBV12 | CASSSVNEQYF | ✔ | ✔ | ✔ | 19, 21 |

B

| Public TCRs | | | No peptide+Ms-IgG-Fab | | | NLV+Ms-IgG-Fab | | | No peptide+Mono-OKT3-Fab | | | NLV+Mono-OKT3-Fab | | |
|---|---|---|---|---|---|---|---|---|---|---|---|---|---|---|
| Donor | TCR V beta gene | CDR3 Amino Acid Sequence | TCR Abundance | Rank of 173 | Rank Percentile | TCR Abundance | Rank of 139 | Rank Percentile | TCR Abundance | Rank of 129 | Rank Percentile | TCR Abundance | Rank of 100 | Rank Percentile |
| 53M | TCRBV27-01*01 | CASSLEGYTEAFF (ref 14, 18, 19, 24) | 340 | 133 | 77% | 4,717 | 18 | 13% | 100 | 119 | 92% | 12,317 | 4 | 4% |
| | TCRBV12 | CASSPGTYGYTF (ref. 19, 24) | 0 | 173 | 100% | 10,945 | 6 | 4% | 458 | 84 | 65% | 6,158 | 6 | 6% |
| | TCRBV12 | CASSIMNEQFF | 0 | 173 | 100% | 7,183 | 12 | 9% | 21,212 | 2 | 2% | 5,276 | 7 | 7% |
| | TCRBV27-01*01 | CASSPIAGYPHEQYF | 0 | 173 | 100% | 10,604 | 7 | 5% | 15,904 | 4 | 3% | 8,197 | 5 | 5% |
| | TCRBV06 | CASSPTTGTGTYGYTF | 25 | 170 | 98% | 19,163 | 2 | 1% | 42,683 | 1 | 1% | 16,740 | 3 | 3% |
| | TCR V beta gene | CDR3 Amino Acid Sequence | TCR Abundance | Rank of 102 | Rank Percentile | TCR Abundance | Rank of 92 | Rank Percentile | TCR Abundance | Rank of 123 | Rank Percentile | TCR Abundance | Rank of 109 | Rank Percentile |
| 28M | TCRBV27-01*01 | CASSPIAGYPHEQYF | 47 | 92 | 90% | 3,779 | 6 | 7% | 7,646 | 3 | 2% | 6,004 | 7 | 6% |
| | TCRBV06 | CASSPTTGTGTYGYTF | 215 | 59 | 58% | 7,965 | 3 | 3% | 19,982 | 1 | 1% | 11,132 | 4 | 4% |
| | TCRBV12 | CASSSVNEQFF | 1,224 | 12 | 12% | 90,590 | 1 | 1% | 10,008 | 2 | 2% | 57,318 | 1 | 1% |
| | TCR V beta gene | CDR3 Amino Acid Sequence | TCR Abundance | Rank of 111 | Rank Percentile | TCR Abundance | Rank of 115 | Rank Percentile | TCR Abundance | Rank of 120 | Rank Percentile | TCR Abundance | Rank of 110 | Rank Percentile |
| 47M | TCRBV27-01*01 | CASSPIAGYPHEQYF | 913 | 19 | 17% | 51,372 | 1 | 1% | 4,238 | 10 | 8% | 34,592 | 4 | 4% |
| | TCRBV06 | CASSPTTGTGTYGYTF | 638 | 31 | 28% | 31,414 | 2 | 2% | 497 | 47 | 39% | 48,671 | 1 | 1% |
| | TCRBV14-01*01 | CASSQEPGNYGYTF | 7,087 | 1 | 1% | 4,922 | 6 | 5% | 20,268 | 4 | 3% | 18,324 | 7 | 6% |
| | TCRBV12 | CASSSAYYGYTF | 128 | 83 | 75% | 23,084 | 3 | 3% | 341 | 63 | 53% | 21,567 | 5 | 5% |
| | TCRBV20 | CSANQGGGNTEAFF | 4,615 | 3 | 3% | 3,636 | 8 | 7% | 54,735 | 1 | 1% | 40,691 | 2 | 2% |
| | TCRBV29-01*01 | CSVTLPQADGRYGYTF | 5,202 | 2 | 2% | 4,512 | 7 | 6% | 44,657 | 2 | 2% | 40,088 | 3 | 3% |

| 0 | 25,000 | 50,000 | 75,000 | 100,000 |
|---|---|---|---|---|

Live cell counts

**Supplemental Table 3. Public TCRs identified in the present study.** (**A**) Among three exog-NLV-bulk-responsive donors, 11 public TCRs were found extending to the top-30 ranked clones. Of these, 9 were shared between all three donors, and 2 were in one donor only but have been previously reported in the literature. (**B**) Public TCR occurrences in top-ten ranks of NLV+Mono-OKT3-Fab combinatorial condition. TCR abundance in each of the four culture conditions per donor is shown.

### Top-10 TCR analysis and commentary

1. Genomic DNA was sequenced on the last day (day 9) of recall assays of PBMCs from each donor, having been cultured +/- exogenous NLV peptide and control Ms-IgG-Fab or Mono-OKT3-Fab.

2. The frequencies of individual TCR clone sequences along with live cell counts from flow cytometry were used to estimate the number of each unique TCR-bearing clone (TCR abundance) at day nine for each sample (see figure to the right).

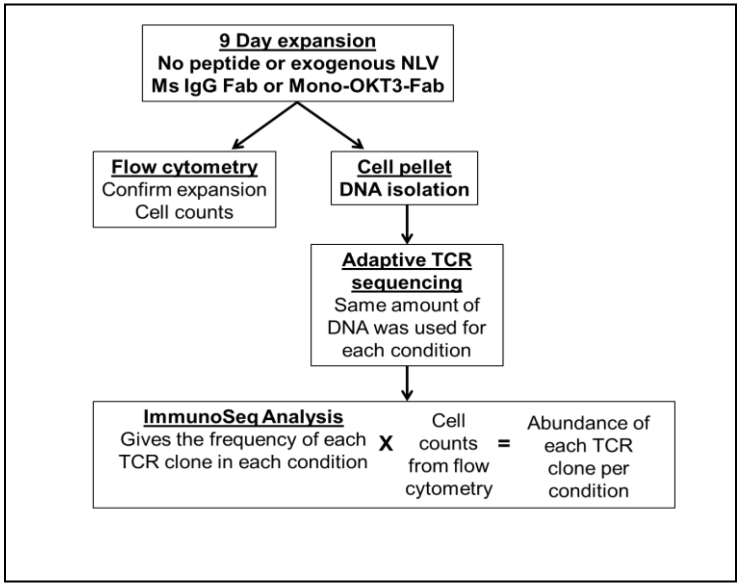


3. TCR clones with the same TCR abundance were equally ranked. (For example, TCR abundances of 9000, 7500, 7500, 3000 would be ranked 1, 2, 2, 3, respectively). At the top of each donor, "Rank of X" shows how many total ranks there were for each particular condition. "Rank percentile" was calculated by taking the TCR clone rank divided by the total ranks for that condition.

4. For each donor, TCR clone sequences were ranked by most abundant to least abundant in the NLV+Mono-OKT3-Fab culture condition, whose top-10 ranks are highlighted in yellow, and the TCR abundance and rank of that same amino acid sequence in the other three conditions are also shown.

5. TCR clones for which the NLV+Ms-IgG-Fab condition > No peptide+Ms-IgG-Fab condition by at least 10% were considered NLV-responsive.

6. Exogenous-NLV-non-responsive clones are in red font. Some TCRs had high relative abundance in the No peptide+Ms-IgG-Fab negative-control condition, and exogenous NLV did not increase cell numbers. These clones are not considered exogenous-NLV-responsive.

7. Some top clones could not be accurately assessed for NLV-responsiveness due to failure to sample the same clones in control or comparator conditions. These clones, with a TCR abundance of 0 in one of the four conditions, are in black font without highlighting.

### Supplemental Table 4. TCR abundance and clone ranks from Exog-NLV-bulk-non-responsive donors.

| ᴐonor | Exog-NLV-bulk-non-responsive donors | | No peptide+Ms-IgG-Fab | | | NLV+Ms-IgG-Fab | | | No peptide+Mono-OKT3-Fab | | | NLV+Mono-OKT3-Fab | | | Notes |
|---|---|---|---|---|---|---|---|---|---|---|---|---|---|---|---|
| 74M | TCR V beta gene | CDR3 Amino Acid Sequence | TCR Abundance | Rank of 122 | Rank Percentile | TCR Abundance | Rank of 107 | Rank Percentile | TCR Abundance | Rank of 131 | Rank Percentile | TCR Abundance | Rank of 104 | Rank Percentile | |
| | TCRBV12 | CASRSLRDLNTEAFF | 611 | 41 | 34% | 774 | 22 | 21% | 13,325 | 2 | 2% | 32,446 | 1 | 1% | |
| | TCRBV05-05*01 | CASSSGANQPQHF | 9,543 | 1 | 1% | 9,749 | 2 | 2% | 16,225 | 1 | 1% | 8,192 | 2 | 2% | <10% increase from No peptide+Ms-IgG-Fab to NLV+Ms-IgG-Fab, so not considered NLV-resposive |
| | TCRBV06 | CASSENRGNSYEQYF | 2,416 | 6 | 5% | 1,346 | 11 | 10% | 6,149 | 5 | 4% | 6,133 | 3 | 3% | |
| | TCRBV27-01*01 | CASSPQTGSSYEQYF | 218 | 83 | 68% | 121 | 76 | 71% | 2,649 | 7 | 5% | 4,250 | 4 | 4% | |
| | TCRBV27-01*01 | CASSPSDTSSYEQYF | 234 | 80 | 66% | 152 | 69 | 64% | 2,609 | 8 | 6% | 3,850 | 5 | 5% | |
| | TCRBV03 | CASSQVPDSDCNQPQHF | 115 | 101 | 83% | 132 | 73 | 68% | 1,093 | 18 | 14% | 2,797 | 6 | 6% | |
| | TCRBV27-01*01 | CASSPQTGTTYEQYF | 147 | 95 | 78% | 78 | 87 | 81% | 1,574 | 11 | 8% | 2,152 | 7 | 7% | |
| | TCRBV02-01*01 | CASSEEWGTSGGANEQFF | 27 | 117 | 96% | 39 | 97 | 91% | 630 | 33 | 25% | 1,660 | 8 | 8% | |
| | TCRBV02-01*01 | CASSEFPGQGDEKLFF | 0 | 122 | 100% | 0 | 107 | 100% | 0 | 131 | 100% | 1,606 | 9 | 9% | |
| | TCRBV24 | CATSEPTGGELFF | 0 | 122 | 100% | 0 | 107 | 100% | 0 | 131 | 100% | 1,606 | 9 | 9% | |
| 78F | TCR V beta gene | CDR3 Amino Acid Sequence | TCR Abundance | Rank of 109 | Rank Percentile | TCR Abundance | Rank of 126 | Rank Percentile | TCR Abundance | Rank of 130 | Rank Percentile | TCR Abundance | Rank of 117 | Rank Percentile | |
| | TCRBV27-01*01 | CASCSTTGYETQYF | 951 | 19 | 17% | 1,467 | 9 | 7% | 5,467 | 3 | 2% | 23,002 | 1 | 1% | |
| | TCRBV19-01 | CASSRVSGSNEQYF | 39,477 | 1 | 1% | 29,505 | 1 | 1% | 72,216 | 1 | 1% | 13,693 | 2 | 2% | |
| | TCRBV03 | CASSQDVGDTEAFF | 158 | 77 | 71% | 142 | 84 | 67% | 2,478 | 13 | 10% | 5,336 | 3 | 3% | |
| | TCRBV20 | CSATGQPYGYTF | 50 | 99 | 91% | 30 | 117 | 93% | 2,485 | 12 | 9% | 4,579 | 4 | 3% | |
| | TCRBV27-01*01 | CASSLGSSPGELFF | 99 | 89 | 82% | 98 | 97 | 77% | 1,575 | 19 | 15% | 4,080 | 5 | 4% | |
| | TCRBV06-05*01 | CASGGAGPYNEQFF | 158 | 77 | 71% | 51 | 111 | 88% | 1,315 | 27 | 21% | 3,261 | 6 | 5% | |
| | TCRBV19-01 | CASSAPGQVLNEKLFF | 198 | 71 | 65% | 162 | 79 | 63% | 1,331 | 26 | 20% | 3,243 | 7 | 6% | |
| | TCRBV06 | CATSPGLEAFF | 89 | 91 | 83% | 94 | 98 | 78% | 833 | 49 | 38% | 2,815 | 8 | 7% | <10% increase from No peptide+Ms-IgG-Fab to NLV+Ms-IgG-Fab, so not considered NLV-resposive |
| | TCRBV07-02*01 | CASSLAETENTEAFF | 589 | 32 | 29% | 678 | 23 | 18% | 2,554 | 11 | 8% | 2,762 | 9 | 8% | |
| | TCRBV09-01 | CASSAQGSKPQHF | 267 | 59 | 54% | 172 | 76 | 60% | 1,728 | 15 | 12% | 2,726 | 10 | 9% | |
| 59F | TCR V beta gene | CDR3 Amino Acid Sequence | TCR Abundance | Rank of 101 | Rank Percentile | TCR Abundance | Rank of 136 | Rank Percentile | TCR Abundance | Rank of 104 | Rank Percentile | TCR Abundance | Rank of 106 | Rank Percentile | |
| | TCRBV07-09 | CASSSRFGTGTHEQYF | 3,990 | 2 | 2% | 4,453 | 3 | 2% | 13,023 | 1 | 1% | 16,143 | 1 | 1% | |
| | TCRBV04-03*01 | CASSQDYPPAGGTNNEQFF | 231 | 62 | 61% | 431 | 53 | 39% | 4,886 | 4 | 4% | 11,334 | 2 | 2% | |
| | TCRBV29-01*01 | CSVEDEDSRTDTQYF | 104 | 83 | 82% | 227 | 88 | 65% | 2,417 | 7 | 7% | 7,824 | 3 | 3% | |
| | TCRBV20 | CSAGRGIKTGRSETQYF | 63 | 90 | 89% | 237 | 86 | 63% | 904 | 28 | 27% | 5,360 | 4 | 4% | |
| | TCRBV10-02*01 | CASSAGTGTYGYTF | 259 | 58 | 57% | 232 | 87 | 64% | 5,399 | 2 | 2% | 4,169 | 5 | 5% | |
| | TCRBV02-01*01 | CASSVRDYEQYF | 306 | 53 | 52% | 317 | 71 | 52% | 3,138 | 6 | 6% | 3,898 | 6 | 6% | <10% increase from No peptide+Ms-IgG-Fab to NLV+Ms-IgG-Fab, so not considered NLV-resposive |
| | TCRBV03 | CASSRQRTYTGELFF | 23 | 97 | 96% | 57 | 124 | 91% | 1,313 | 21 | 20% | 3,537 | 7 | 7% | |
| | TCRBV06-05*01 | CASSVAGGLQETQYF | 17 | 98 | 97% | 66 | 122 | 90% | 261 | 75 | 72% | 3,050 | 8 | 8% | |
| | TCRBV11-02*02 | CASSLVGVEAFF | 52 | 92 | 91% | 62 | 123 | 90% | 1,226 | 24 | 23% | 2,987 | 9 | 8% | |
| | TCRBV07-09 | CASSLQTGVAFF | 986 | 20 | 20% | 1,760 | 15 | 11% | 1,539 | 16 | 15% | 2,274 | 10 | 9% | |

**Supplemental Table 4. TCR abundance and clone ranks from Exog-NLV-bulk-non-responsive donors.**

**Exog-NLV-bulk-responsive Donors**

| Donor | TCR Vβ Gene | Amino Acid Sequence | TCR Abundance (Exog NLV: -; Fab: Ms-IgG) | Rank (Exog NLV: -; Fab: Ms-IgG) | TCR Abundance (Exog NLV: +; Fab: Ms-IgG) | Rank (Exog NLV: +; Fab: Ms-IgG) | TCR Abundance (Exog NLV: -; Fab: Mono-OKT3) | Rank (Exog NLV: -; Fab: Mono-OKT3) | TCR Abundance (Exog NLV: +; Fab: Mono-OKT3) | Rank (Exog NLV: +; Fab: Mono-OKT3) | Fold change of live T cell abundance: NLV+Ms-IgG-Fab / No peptide+Ms-IgG-Fab | Fold change of live T cell abundance: NLV+Mono-OKT3-Fab / No peptide+Mono-OKT3-Fab | Fold change of live T cell abundance: NLV+Mono-OKT3-Fab / NLV+Ms-IgG-Fab |
|---|---|---|---|---|---|---|---|---|---|---|---|---|---|
| 72F | TCRBV06 | CASSKVTGTGNYGYTF | 1,511 | 26 | 130,547 | 1 | 125,632 | 1 | 214,687 | 1 | 86.42 | 1.71 | 1.64 |
| | TCRBV06-05*01 | CASSPSTGTIYGYTF | 118 | 109 | 8,254 | 3 | 7,241 | 4 | 19,561 | 3 | 70.20 | 2.70 | 2.37 |
| | TCRBV06 | CASSPITGQGAYGYTF | 428 | 60 | 4,163 | 5 | 6,263 | 5 | 8,800 | 4 | 9.74 | 1.41 | 2.11 |
| | TCRBV11-02*02 | CASSPWAYATDTQYF | 29 | 134 | 1,284 | 15 | 619 | 39 | 6,652 | 6 | 45.04 | 10.74 | 5.18 |
| 53M | TCRBV05-08*01 | CASSRTSINEQFF | 110 | 160 | 52,021 | 1 | 627 | 67 | 90,986 | 1 | 470.82 | 145.02 | 1.75 |
| | **TCRBV27-01*01** | **CASSLEGYTEAFF** | 340 | 133 | 4,717 | 18 | 100 | 119 | 12,317 | 4 | 13.87 | 123.68 | 2.61 |
| 28M | TCRBV06 | CASTPQTGTGYYGYTF | 294 | 48 | 7,420 | 5 | 160 | 86 | 12,250 | 3 | 25.22 | 76.56 | 1.65 |
| 47M | **TCRBV06** | **CASSPTTGTGTYGYTF** | 638 | 31 | 31,414 | 2 | 497 | 47 | 48,671 | 1 | 49.27 | 97.93 | 1.55 |

**Exog-NLV-bulk-non-responsive Donors**

| Donor | TCR Vβ Gene | Amino Acid Sequence | TCR Abundance (Exog NLV: -; Fab: Ms-IgG) | Rank (Exog NLV: -; Fab: Ms-IgG) | TCR Abundance (Exog NLV: +; Fab: Ms-IgG) | Rank (Exog NLV: +; Fab: Ms-IgG) | TCR Abundance (Exog NLV: -; Fab: Mono-OKT3) | Rank (Exog NLV: -; Fab: Mono-OKT3) | TCR Abundance (Exog NLV: +; Fab: Mono-OKT3) | Rank (Exog NLV: +; Fab: Mono-OKT3) | Fold change of live T cell abundance: NLV+Ms-IgG-Fab / No peptide+Ms-IgG-Fab | Fold change of live T cell abundance: NLV+Mono-OKT3-Fab / No peptide+Mono-OKT3-Fab | Fold change of live T cell abundance: NLV+Mono-OKT3-Fab / NLV+Ms-IgG-Fab |
|---|---|---|---|---|---|---|---|---|---|---|---|---|---|
| **74M** | TCRBV12 | CASRSLRDLNTEAFF | 611 | 41 | 774 | 22 | 13,325 | 2 | 32,446 | 1 | 1.27 | 2.44 | 41.91 |
| | TCRBV03 | CASSQVPDSDCNQPQHF | 115 | 101 | 132 | 73 | 1,093 | 18 | 2,797 | 6 | 1.16 | 2.56 | 21.15 |
| | TCRBV02-01*01 | CASSEEWGTSGGANEQFF | 27 | 117 | 39 | 97 | 630 | 33 | 1,660 | 8 | 1.43 | 2.63 | 42.67 |
| **78F** | TCRBV27-01*01 | CASCSTTGYETQYF | 951 | 19 | 1,467 | 9 | 5,467 | 3 | 23,002 | 1 | 1.54 | 4.21 | 15.68 |
| | TCRBV07-02*01 | CASSLAETENTEAFF | 589 | 32 | 678 | 23 | 2,554 | 11 | 2,762 | 9 | 1.15 | 1.08 | 4.08 |
| **59F** | TCRBV07-09 | CASSSRFGTGTHEQYF | 3,990 | 2 | 4,453 | 3 | 13,023 | 1 | 16,143 | 1 | 1.12 | 1.24 | 3.63 |
| | TCRBV04-03*01 | CASSQDYPPAGGTNNEQFF | 231 | 62 | 431 | 53 | 4,886 | 4 | 11,334 | 2 | 1.87 | 2.32 | 26.32 |
| | TCRBV29-01*01 | CSVEDEDSRTDTQYF | 104 | 83 | 227 | 88 | 2,417 | 7 | 7,824 | 3 | 2.19 | 3.24 | 34.45 |
| | TCRBV20 | CSAGRGIKTGRSETQYF | 63 | 90 | 237 | 86 | 904 | 28 | 5,360 | 4 | 3.73 | 5.93 | 22.66 |
| | TCRBV03 | CASSRQRTYTGELFF | 23 | 97 | 57 | 124 | 1,313 | 21 | 3,537 | 7 | 2.46 | 2.69 | 62.30 |
| | TCRBV06-05*01 | CASSVAGGLQETQYF | 17 | 98 | 66 | 122 | 261 | 75 | 3,050 | 8 | 3.83 | 11.69 | 46.04 |
| | TCRBV11-02*02 | CASSLVGVEAFF | 52 | 92 | 62 | 123 | 1,226 | 24 | 2,987 | 9 | 1.19 | 2.44 | 48.56 |
| | TCRBV07-09 | CASSLQTGVAFF | 986 | 20 | 1,760 | 15 | 1,539 | 16 | 2,274 | 10 | 1.79 | 1.48 | 1.29 |

**Supplemental Table 5. Fold changes in TCR abundance of "Gold" response clones.** "Gold" awarded TCR clones ranked in the top-10 of NLV+Mono-OKT3-Fab condition as described in Table 1 and Figure 5B were analyzed for fold changes of live T cell abundance between various conditions in exog-NLV-bulk-responsive and exog-NLV-bulk-non-responsive donors. Bold sequences indicate public TCRs.

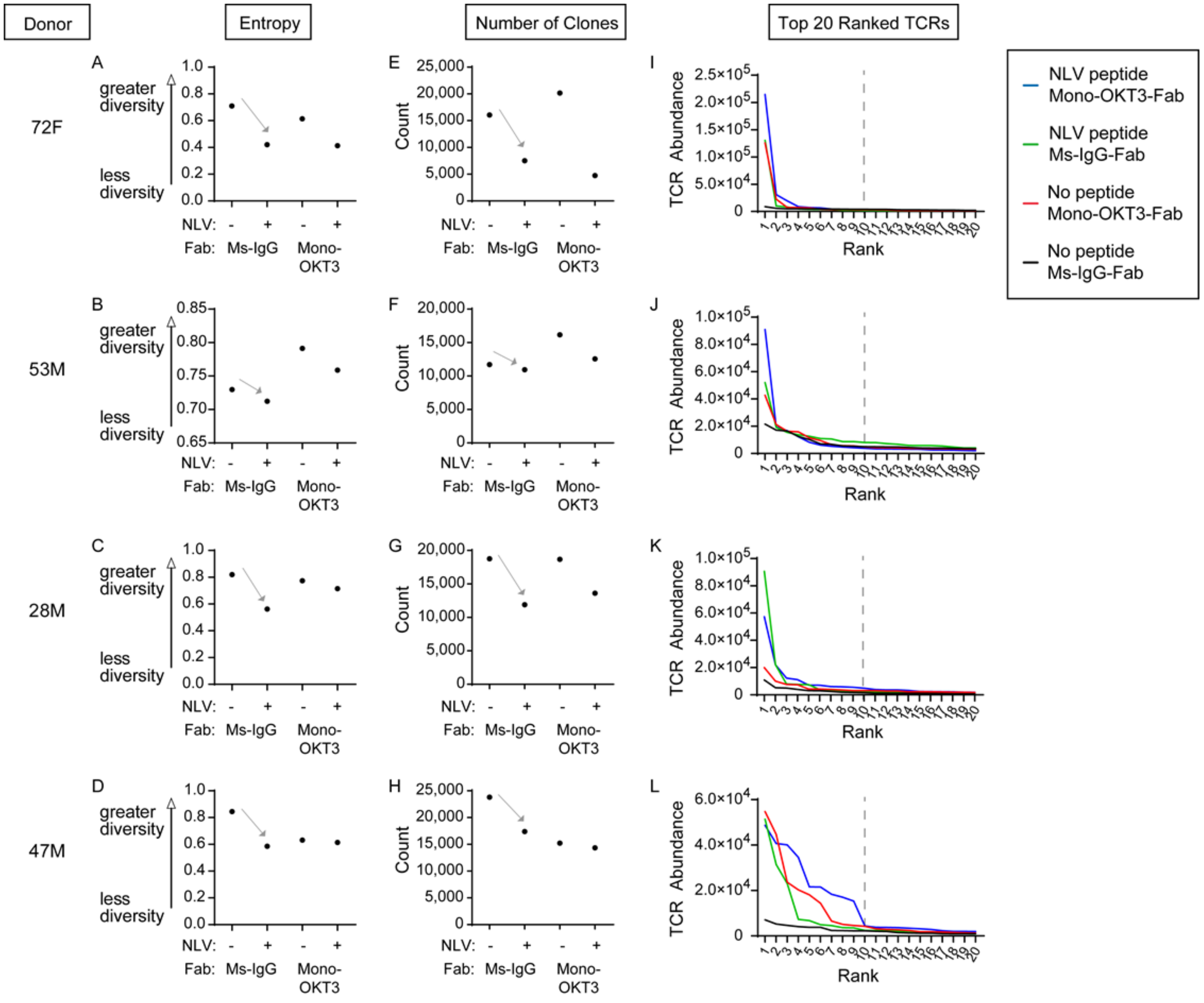


**Supplemental Figure 1. TCR repertoire complexity in recall assays from exog-NLV-bulk-responsive donors.** (**A-H**) For each donor, the four culture conditions were evaluated for clonal diversity by calculating scaled Shannon entropy and counting unique TRBV-CDR3-bearing T cell clones in sampled DNA. Exogenous NLV drove a decrease in both of these parameters. (**I-L**) The number of live T cells bearing each clone was estimated using single-cell ImmunoSeq data and total live T cell counts for whole samples. Live clonal abundances were ranked from highest to lowest for each culture condition and donor. Major differences in rank-versus-rank performance between conditions were concentrated in the top-10 clones.

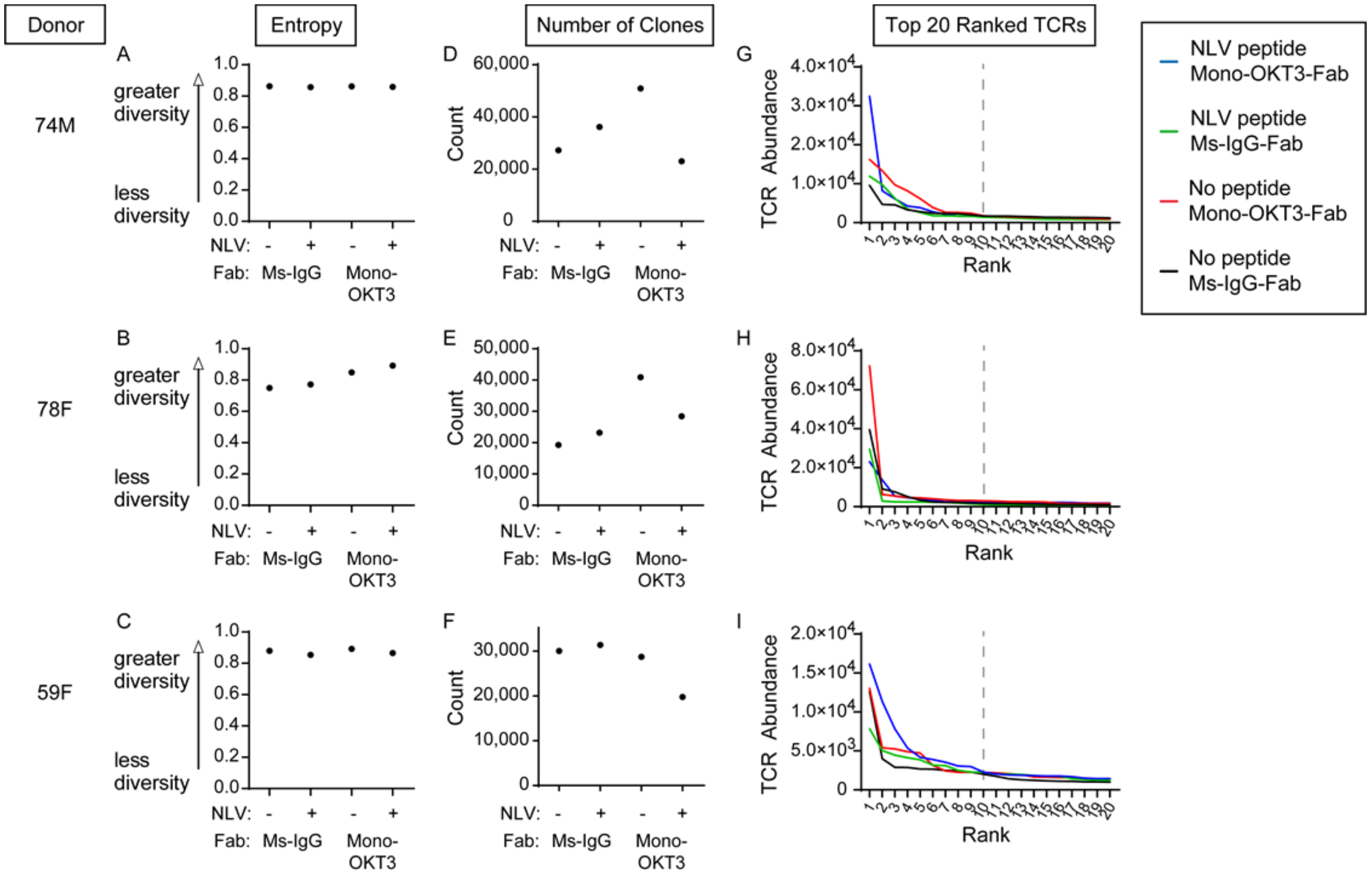


**Supplemental Figure 2. TCR repertoire complexity in recall assays from exog-NLV-bulk-non-responsive donors.** (**A-F**) For each donor, the four culture conditions were evaluated for clonal diversity by calculating scaled Shannon entropy and counting unique TRBV-CDR3-bearing T cell clones in sampled DNA. (**G-I**) The number of live T cells bearing each clone was estimated using single-cell ImmunoSeq data and total live T cell counts for whole samples. Live clonal abundances were ranked from highest to lowest for each culture condition and donor. Major differences in rank-versus-rank performance between conditions were concentrated in the top-10 clones.
