## Supplementary figures and images for "Public and private human T cell clones respond differentially to HCMV antigen when boosted by CD3 co-potentiation"

### Graphical abstract

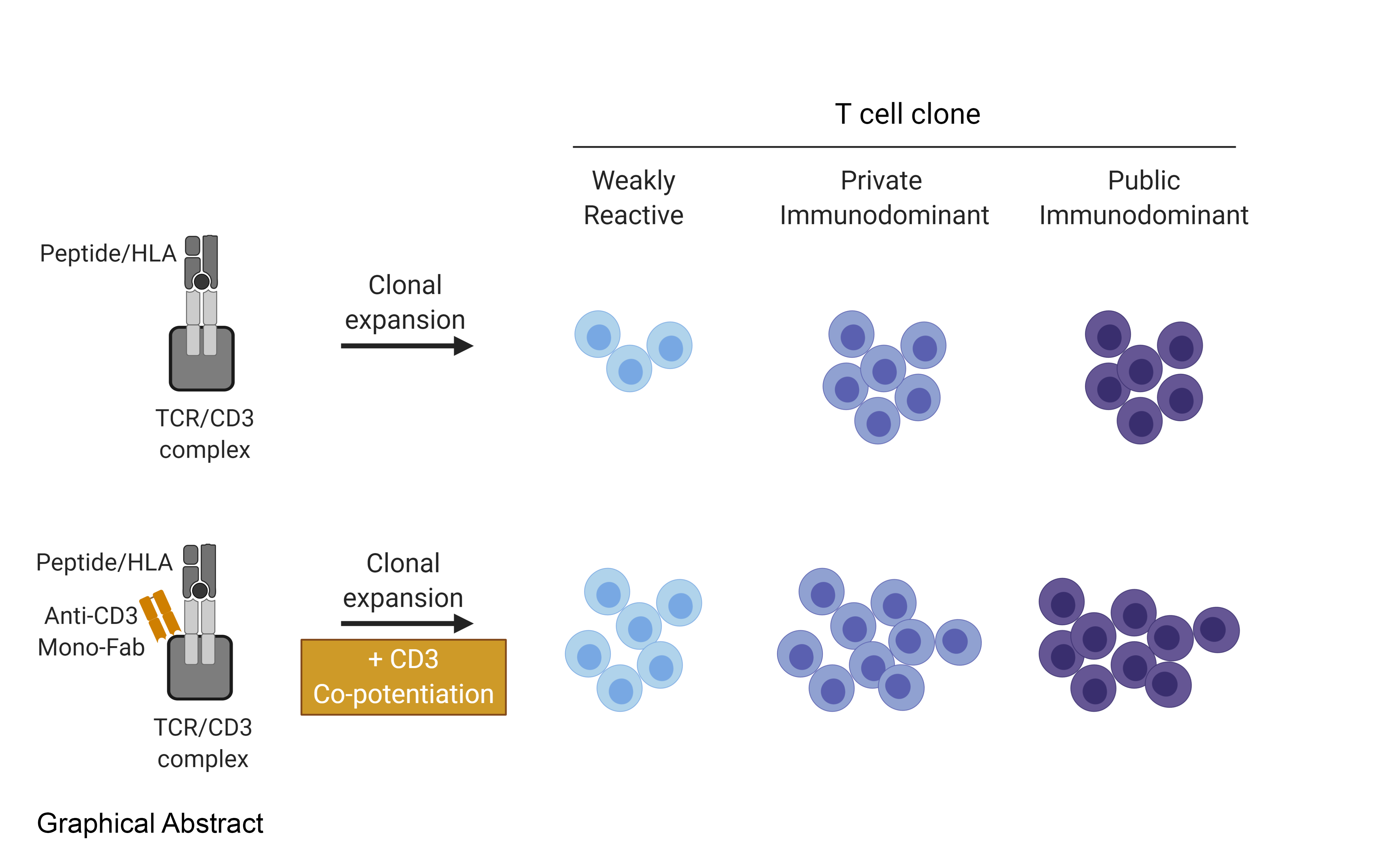
